# Plasma membrane mediated GLUT10 mitochondrial targeting regulates intracellular ascorbic acid homeostasis

**DOI:** 10.1101/2025.01.09.632083

**Authors:** Anu Chirackal Jose, Yu-Wei Syu, Hao-Wen Lai, Ming-Yuan Tsai, Yi-Fan Jiang, Shao-Chun Hsu, Po-Yen Lin, Wan-Chen Huang, Wei-Chen Chu, Chi-Yu Fu, Yi-Ching Lee

## Abstract

Regulation of intracellular ascorbic acid (AA) levels is critical for connective tissue development and maintenance, but the underlying mechanisms remain unclear. In this study, we uncover a novel regulatory role of glucose transporter 10 (GLUT10) in AA homeostasis through a noncanonical trafficking pathway from the endomembrane system to mitochondria, traditionally considered as separate entities. We demonstrate that GLUT10 transit from ER to mitochondria through the plasma membrane (PM) and endosomes increases under stress conditions. This PM localization enhances the transport of dehydroascorbic acid (DHA), the oxidized form of AA, thereby maintaining intracellular AA levels. Disruption of this trafficking impairs AA homeostasis. Our findings reveal a previously unrecognized localization of GLUT10 at the PM and endosomes, highlighting a novel communication between endomembrane system and mitochondria for proetin redistribution, which is essential for maintaining intracellular AA homeostasis and facilitating environmental adaptation.

**Graphical abstract:** 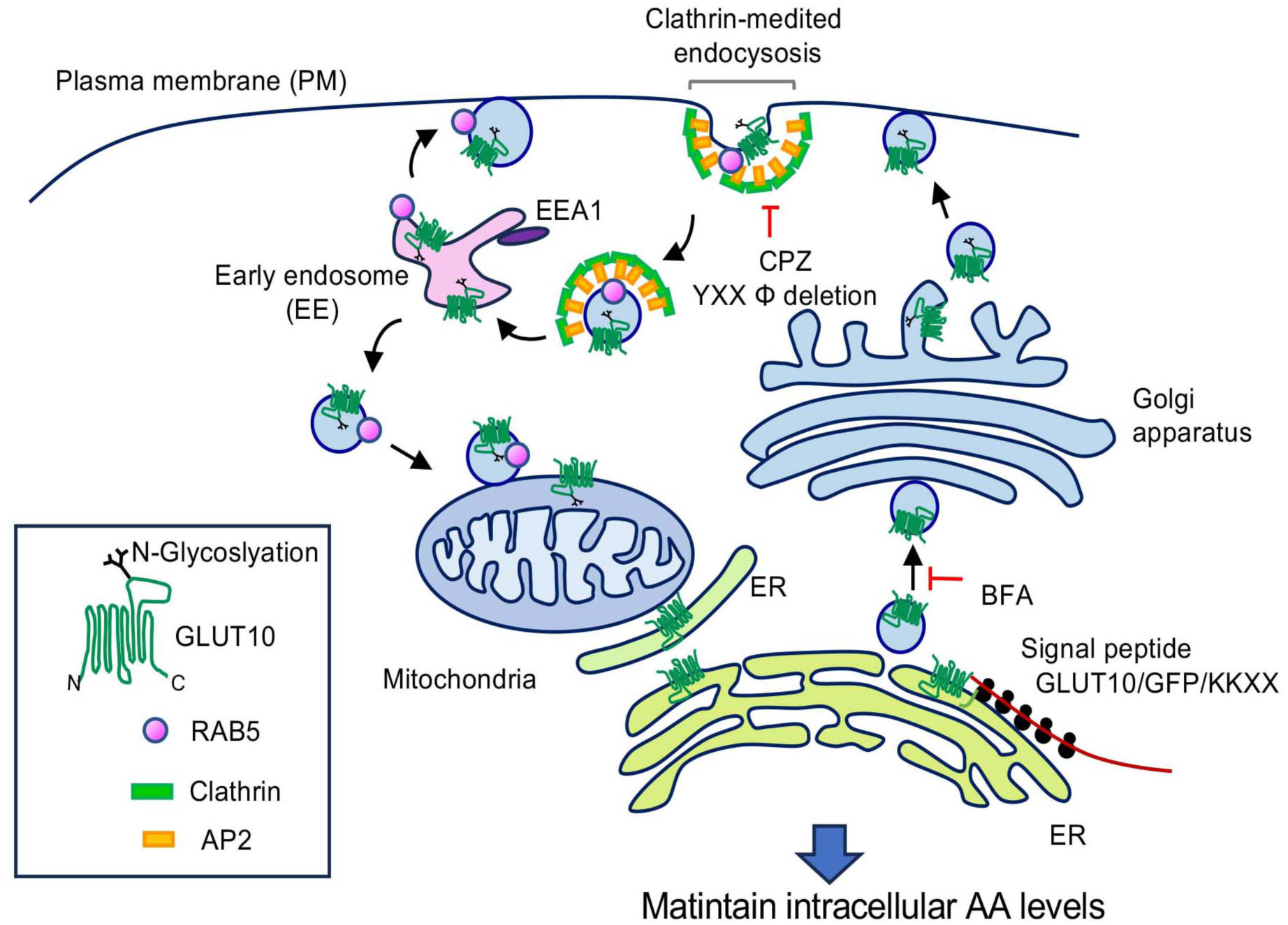

## Introduction

For a protein to function optimally, it is often necessary to establish proper physiological context by directing it to an appropriate subcellular localization. Many proteins play functional roles in cellular compartments distant from their location of synthesis, and targeting the protein to its proper setting typically involves intricate targeting mechanisms that are dependent on sorting signals in the amino acid sequence ^1^. It is widely accepted that the mechanisms targeting proteins to mitochondria are largely distinct from those targeting other proteins to the endomembrane system. In general, nuclear encoded mitochondrial proteins are synthesized in the cytosol and rely on mitochondrial targeting sequences to enter the mitochondrial compartments. Meanwhile, proteins destined for secretion or the endomembrane system [i.e., endoplasmic reticulum (ER), Golgi, lysosomes and plasma membrane (PM)] are synthesized on ribosomes associated with the rough ER ^2^. Intriguingly, recent studies have shown that some proteins can be dually targeted to mitochondria and other compartments ^3^, and several nuclear encoded mitochondrial proteins are N-glycosylated ^4–6^, a process that typically occurs in the ER and Golgi ^7^. Thus, targeting of proteins to mitochondria may be more complex and varied than previously expected.

It is well accepted that mitochondria have many functions and communicate with other organelles to maintain cellular homeostasis. For instance, direct interactions between mitochondria and organelles such as the ER, nucleus and peroxisomes have been implicated in the regulation of calcium homeostasis and lipid metabolism ^8^, and the interaction and crosstalk between ER and mitochondria have recently garnered significant attention ^9^. The interactions between endosomes and mitochondria have also been shown to facilitate intracellular iron transfer in different types of cells ^10,11^. Although intricate networks of inter-organelle communication that control cellular homeostasis and adaptation are beginning to be identified, the complete landscape of functional organelle interactions remains to be fully delineated.

Proteins of the solute carrier transporter family (SLCs) are structurally characterized by multiple transmembrane domains and function to mediate metabolite movement across all cellular membranes. As such, proper subcellular localization of SLCs is crucial for regulating metabolic flux and maintaining cellular physiological states. However, the mechanisms by which environmental cues regulate the subcellular localization of most SLCs to maintain metabolic homeostasis remain largely unknown. Functional deficiencies in various SLCs have been linked to the etiology of numerous diseases ^12^, providing a valuable platform for studying their function and regulation. Loss-of-function mutations in *SLC2A10* gene, which encodes glucose transporter 10 (GLUT10), lead to arterial tortuosity syndrome (ATS), a rare connective tissue disease that affects the development and maintenance of connective tissues ^13^. Our studies have shown that GLUT10 is highly expressed in aortic smooth muscle cells (ASMCs) of major arteries, the primary site of lethal complications in ATS patients ^14^. In ASMCs, GLUT10 is located in both ER and mitochondria, facilitating the transport of dehydroascorbic acid (DHA), the oxidized form of ascorbic acid (AA; vitamin C) ^14–16^. Overexpression of GLUT10 in ASMCs, significantly increased DHA uptake, maintaining intracellular and mitochondrial AA levels, redox balance and mitochondrial function. Furthermore, AA serves as an enzyme cofactor in the hydroxylation of proline and lysine, which is essential for stabilizing collagen structure within connective tissues. Thus, defects in GLUT10 function impair connective tissue development and integrity, driving ATS pathogenesis ^14,15^.

Intriguingly, our previous work has shown that under oxidative stress and aging conditions, increased GLUT10 colocalization with mitochondria leads to enhanced DHA uptake and elevated intracellular AA levels ^15^, highlighting the importance of its proper localization. This raises a key question: how does the GLUT10 traffic to mitochondria and influence its function, particularly under stress conditions? Our findings reveal a surprising trafficking pathway that directs GLUT10 from ER to mitochondria through PM and endosomal mediated pathway. GLUT10 became N-glycosylated in ER and Golgi then targeting to PM, the N-glycosylation may facilitate early endosomal sorting and subsequent targeting to mitochondria. Furthermore, stress conditions enhance GLUT10 trafficking through the PM and endosomal mitochondrial targeting, enhancing DHA uptake and maintaining intracellular AA levels. This pathway uncovers a previously unrecognized GLUT10 intracellular localization to PM and endosomal vesicles, this is important for the redistribution of GLUT10 to PM and mitochondria for cellular adaptation. Overall, our study establishes a link between protein subcellular localization and the regulation of intracellular AA levels, highlighting the importance of this mechanism for cellular adaptation to environmental stress.

## Results

### GLUT10 transit from the ER to mitochondria via vesicles

To understand how the subcellular localization of GLUT10 in ER and mitochondria may change under conditions of oxidative stress, we utilized a live-cell imaging system to observe GLUT10 intracellular trafficking in A10 rat ASMCs expressing GLUT10/GFP and Mito/DsRed (Fig. 1A). Initially, GLUT10 was localized to the perinuclear region (i.e., ER), but addition of H_2_O_2_ resulted in a whole-cell redistribution of GLUT10, which involved a gradual increase in mitochondrial colocalization (Fig. 1B and C; Movie S1). By capturing live-cell images at 1-s intervals, we were able to observe active movement of GLUT10-containing vesicles, which approached, made direct contact with, and merged with mitochondria (Fig. 1D-F; Movie S2 and Movie S3).

**Figure 1.**
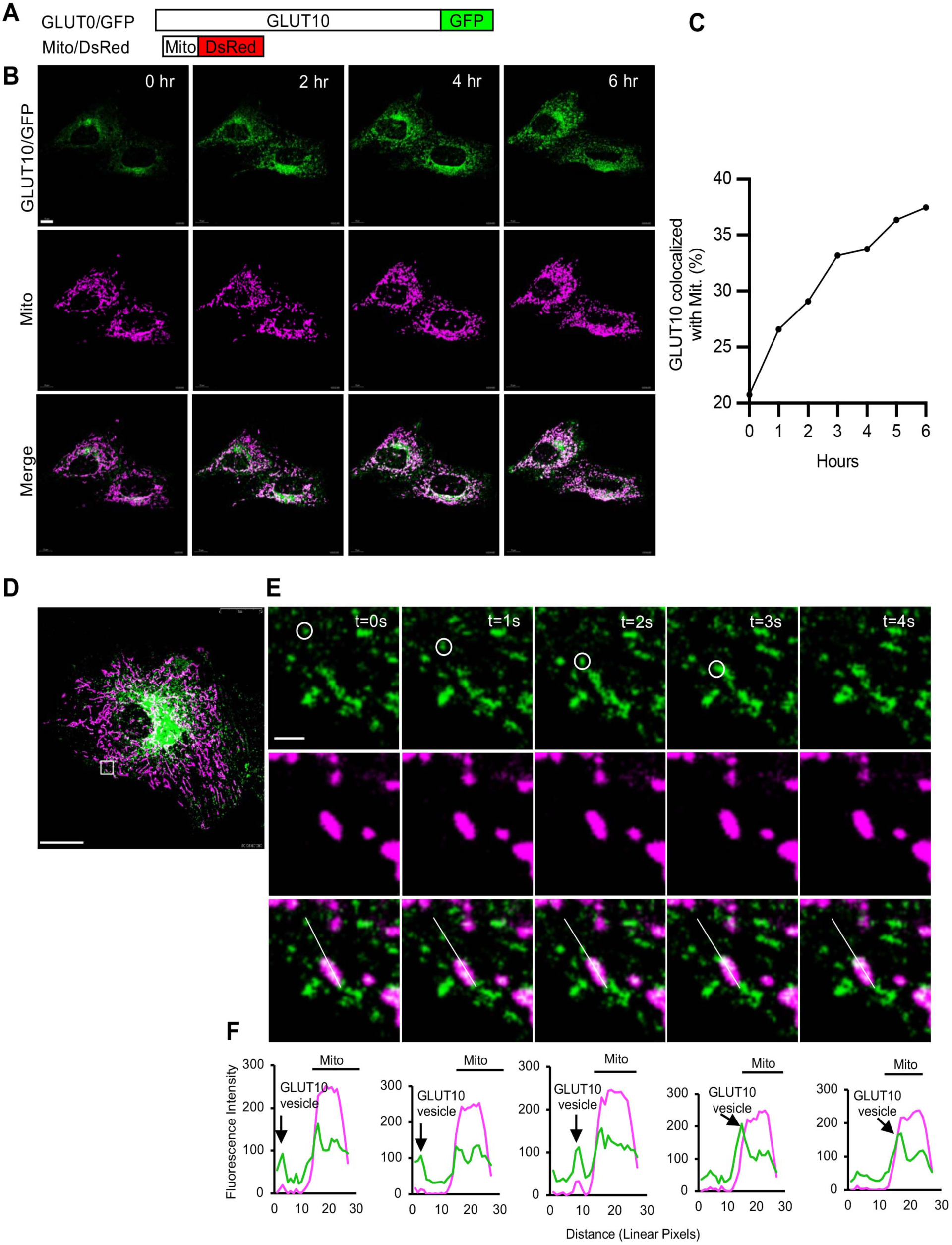
GLUT10-containing vesicles target to mitochondria. (**A**) GLUT10/GFP and Mito/DsRed fusion proteins. (**B**) Confocal images show colocalization of GLUT10/GFP and mitochondria in live cells. A10 cells expressing both GLUT10/GFP and Mito/DsRed were treated with 100 μM H_2_O_2_; imaging was performed every 1 h for 6 h. Scale bar, 10 μm. (**C**) Quantification of the percentage of GLUT10/GFP colocalized with Mito/DsRed, as in **B**. (**D**) Merged confocal images of a live A10 cells expressing GLUT10/GFP and Mito/DsRed. Scale bar, 25 μm. (**E**) Magnified time-lapse confocal images from (**D**); time points from 0-4 s. Scale bar, 1μm. (**F**) Intensity plots along the line from **E** (t = 0-4 s). **B**, **D** and **E**, *Green,* GLUT10/GFP; *magenta*, Mito/DsRed; *white,* merged.

We next wanted to examine the dynamics of GLUT10-containing vesicle targeting to mitochondria, so we analyzed the GLUT10-containing particle movements in H_2_O_2_-treated cells. The particles were tracked at 1-s intervals for 30 s, and we observed linear trajectories of GLUT10 vesicles departing from the perinuclear region toward the cell periphery (Fig. S1A). This pattern of movement was indicative of regulated GLUT10 vesicle trafficking. Our overall analysis of 429 trajectories allowed us to identify two types of motion, including stationary (320 trajectories with track displacements < 0.67 µm) and dynamic (109 trajectories with displacements > 0.67 µm) (Fig. S1B and C). Speed analysis showed that the GLUT10 vesicles slowed upon encountering mitochondria and then merged with stationary tracks (Fig. S1D and E). These observations led us to conclude that the GLUT10-containing vesicles come into close contact with mitochondria, which suggested the vesicles may play a role in targeting GLUT10 to mitochondrial compartments.

### GLUT10 localizes to the endomembrane system and mitochondria

We previously observed GLUT10 localization in both ER and mitochondria of ASMCs ^14,15^. Since our initial live-cell trafficking data revealed a broad intracellular distribution of GLUT10 and an apparent vesicular trafficking process mediating its distribution, we sought to further investigate this phenomenon by thoroughly examining the intracellular distribution of GLUT10. Consistent with previous findings, our confocal imaging experiments on GLUT10/GFP-expressing rate ASMCs, MOVAS cells, showed that the protein colocalized with both ER and mitochondria (Fig. S2A and B). This result was then validated by subcellular fractionation experiments (Fig. S2C and D). Importantly, we further confirmed that endogenous GLUT10 is present in both the ER and mitochondria of human ASMCs and 293T cells (Fig. S2 E-G). Consequently, both ASMCs and 293T cells were used in subsequent experiments.

To explore the ultrastructural details of GLUT10 localization, we performed transmission electron microscopy (TEM) with immunogold labeling. Gold particles were observed on the ER, mitochondria, PM, nuclear envelope (NE), Golgi complex and vesicle structures, with high signals associated with vesicles and ER (Fig. 2A-F). The high density of gold particles on intracellular membrane structures corroborated the localization patterns observed in confocal images. To confirm these results using another experimental approach, we utilized cells expressing enhanced ascorbate peroxidase 2 (APEX2)-tagged GLUT10 (GLUT10/APEX2) and observed the localization of the fusion protein by TEM. Since APEX2 catalyzes diaminobenzidine (DAB) polymerization to provide EM contrast ^17^, we could directly visualize GLUT10/APEX2 subcellular localization (Fig. S3A-C). Compared to controls with standard osmium fixation (Fig. S3D), APEX2-only controls (Fig. S3E) and negative controls without DAB staining (Fig. S3F), specific DAB-stained GLUT10/APEX2 signals were detected on the ER, PM, endocytosed vesicles and mitochondria (Fig. 2G-L; Movie S4).

**Figure 2.**
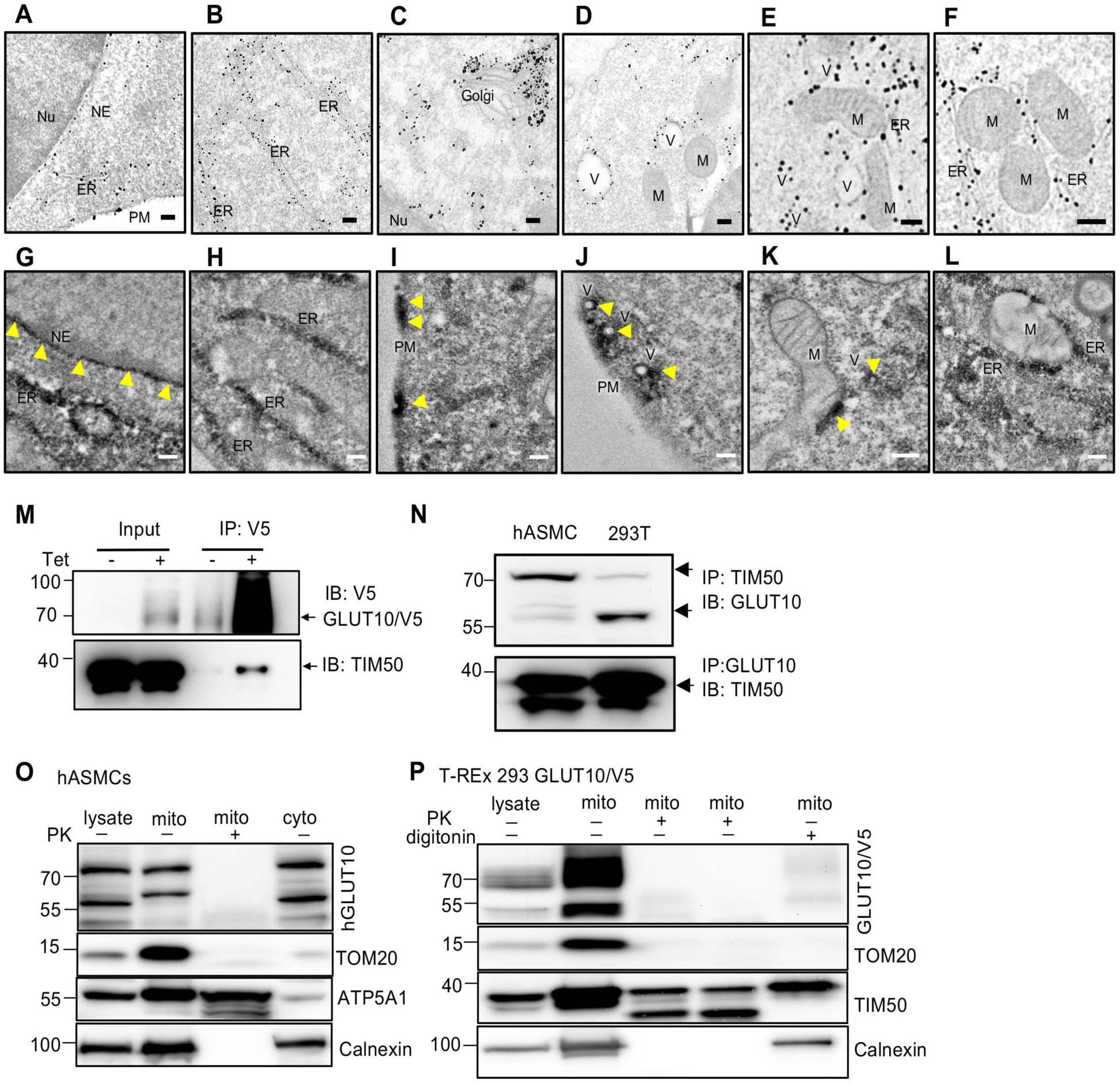
GLUT10 is localized to the endomembrane system and mitochondria. (**A**-**F**) EM micrographs of thin sections of GLUT10/GFP-expressing 293T cells. GFP was labeled by 1.4 nm nanogold clusters. GLUT10/GFP signal was detected in ER (**A, B, E** and **F**), plasma membrane (PM) and nuclear envelope (NE) (**A**), Golgi (**C**), mitochondria (M) (**D-F**), vesicles (V) (**D** and **E**), and ER-mitochondria contacts (**F**). Scale bar, 200 nm. (**G-L**) EM images of MOVAS cells expressing GLUT10/APEX2 show strong EM contrast signals indicating localization of GLUT10/APEX2 in ER (**G, H** and **L**), nuclear envelope (NE) (**G**), plasma membrane (PM) (**I**), vesicles (V) (**J**), mitochondria (**K**), and ER-mitochondria contacts (**L**). Scale bars, 100 nm. (**M and N**) Direct interaction of GLUT10 and TIM50 detected by immunoprecipitation (IP). (**M**) Immunoblot to detect GLUT10/V5 and TIM50 of cell lysates from T-REx-293 cells with or without induction of GLUT10/V5 expression before IP (Input) or after IP with V5. (**N**) Immunoblot detection of GLUT10 and TIM50 using cell lysates from hASMCs and 293T cells; IP with GLUT10 or TIM50. (**O and P**) Localization of GLUT10 on the mitochondrial outer membrane (MOM) examined by immunoblots coupled with proteinase K or digitonin treatment on isolated mitochondria fraction. (**O**) Levels of GLUT10, ATP5A1(mitochondrial inner membrane, MIM), TOM20 (MOM), and Calnexin (ER membrane) in total lysate, cytosolic fractions (cyto), and mitochondrial fractions isolated from hASMCs with or without proteinase K (PK) treatment. (**P**) Levels of GLUT10/V5, TIM50, TOM20 and Calnexin in total lysate and mitochondrial fractions isolated from GLUT10/V5-expressing T-REx 293 cells with or without proteinase K (PK) or digitonin treatment.

Next, we wanted to better understand the mitochondrial localization of GLUT10, so we examined its interactions with mitochondrial proteins. To identify GLUT10-interacting proteins, we conducted co-immunoprecipitation (co-IP) followed by mass spectrometry (MS) on GLUT10/V5-expressing T-REx-293 cells. Using this strategy, we were able to identify several mitochondrial proteins that interact with GLUT10 (Fig. S3A and Table S1), such as import receptor subunits (TOM40, TOM70, TIM50), mitochondrial ATP synthase subunits (ATP5C1, ATP5A1 and ATPAF1), and mitochondrial matrix proteins [NAD (+)-dependent isocitrate dehydrogenases (IDH3B)]. Among these interacting proteins, the most abundant mitochondrial protein interacting with GLUT10/V5 was the mitochondrial inner membrane protein, TIM50 (Fig. S4A). The direct interaction between TIM50 and GLUT10/V5 was further confirmed by co-IP and western blot (Fig. 2M). Additionally, the direct interaction between endogenous GLUT10 and TIM50 was validated in hASMCs and 293T cells by performing co-IP of GLUT10 with TIM50 and reciprocal co-IP of TIM50 with GLUT10 (Fig. 2N). We also explored the potential direct interaction between GLUT10/V5 and a mitochondrial matrix protein, IDH3B, which was identified from the initial co-IP/MS assay. However, no direct interaction was observed for these two proteins (Fig. S4B), underscoring the specificity of the identified direct interaction of GLUT10 with TIM50 and further supporting the conclusion that GLUT10 is indeed localized to mitochondria.

To clarify whether GLUT10 localizes to the mitochondrial outer membrane or the mitochondrial inner membrane, both of which could potentially allow interaction with TIM50, we probed the sub-mitochondrial localization of GLUT10 using a proteinase K and digitonin protection assay ^18^. Mitochondria purified from hASMCs or GLUT10/V5-expressing T-REx-293 cells were subjected to proteinase K digestion, which degraded the cytosol-exposed outer membrane proteins. As expected, the known outer membrane protein TOM20 was sensitive to proteinase K digestion, while inner membrane proteins ATP5A1 and TIM50 were resistant (Fig. 2O and P). Interestingly, GLUT10 was sensitive to proteinase K treatment, suggesting it is most likely localized to the outer membrane (Fig. 2O and P). Next, we wanted to confirm the localization of GLUT10 to the outer membrane and exclude the potential interference of ER contamination on the proteinase K assay results. To do so, mitochondria were purified from cells and treated with digitonin at a concentration that selectively solubilizes the mitochondrial outer membrane but preserves the integrity of the inner membrane and ER (Fig. 2P). After digitonin treatment, both TOM20 and GLUT10 levels were significantly reduced, whereas the inner membrane protein TIM50 and ER membrane protein Calnexin remained unaffected (Fig. 2P), reinforcing the conclusion that GLUT10 is localized to the mitochondrial outer membrane. Taken together, our findings to this point indicated that GLUT10 is localized to both the endomembrane system and the outer membrane of mitochondria.

### ER-Golgi derived N-glycosylated GLUT10 observed in mitochondria

To identify the trafficking pathways that regulate GLUT10 intracellular distribution, we searched for functional targeting motifs within the GLUT10 sequence. By doing so, we identified a typical evolutionarily conserved N-terminal signal peptide (SP) for ER targeting (Fig. 3A and B); however, no consensus mitochondrial targeting signal was found (Fig. 3A). To test the functionality of the SP, we fused the SP of GLUT10 to a GFP tagged with an ER-retention signal (KDEL) (SP/GFP/KDEL) (Fig. 3C). The SP directed GFP to the ER, and the KDEL signal restrained the protein in the ER (Fig. 3D and F), confirming that the SP of GLUT10 is functional and sufficient for ER targeting.

**Figure 3.**
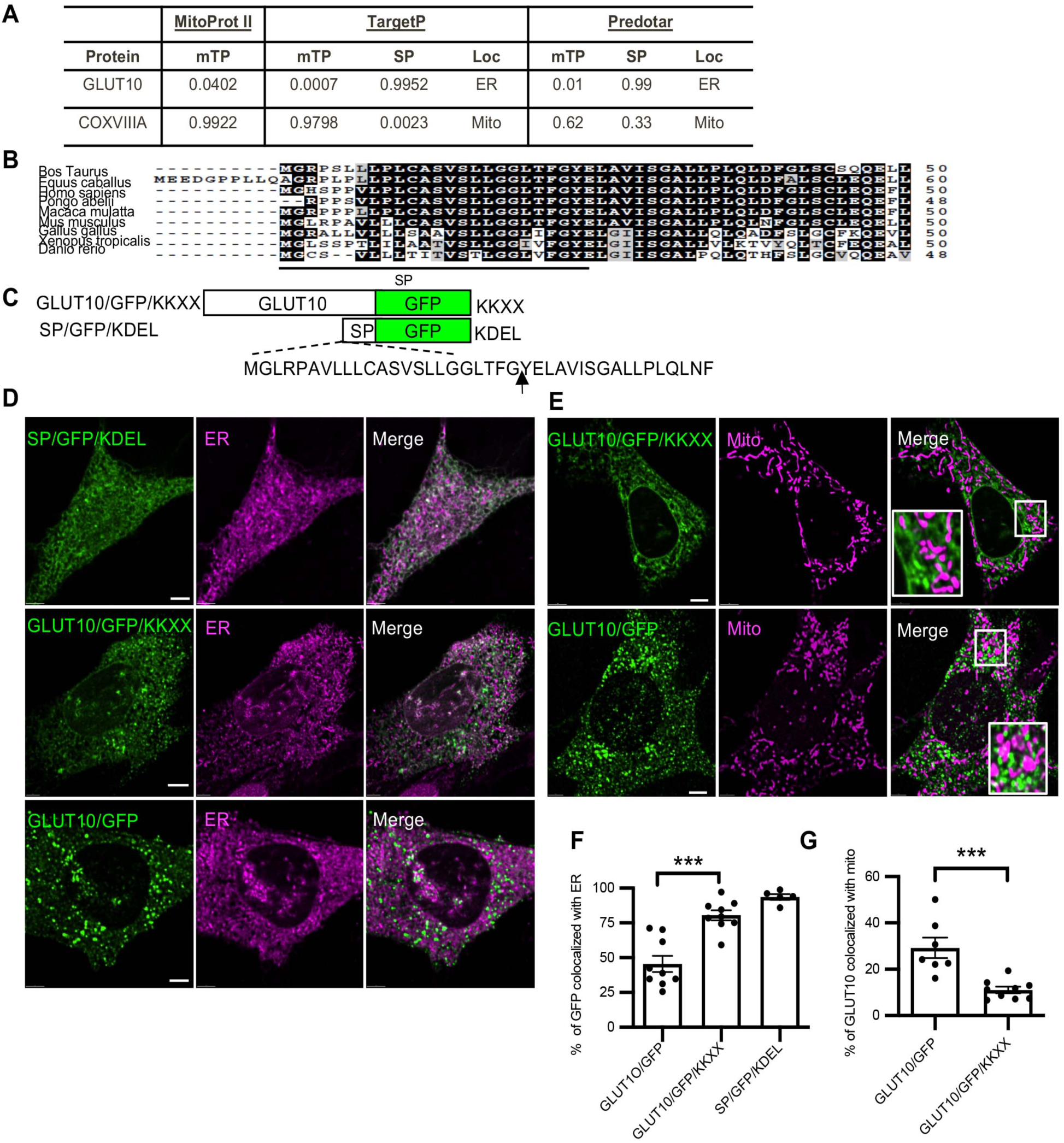
GLUT10 retention in ER reduces its mitochondrial targeting. (**A**) Prediction of mitochondrial-targeting signals (mTP), signal peptide (SP), and subcellular localization (Loc) of mouse GLUT10 by Mitoprot II, TargetP, and Predotar analyses. CoxVIIIA, human cytochrome oxidase subunit VIIIA, a mitochondrial transmembrane protein. (**B**) N-terminal amino acid alignments of GLUT10 from several species was performed with the CLUSTAL W program. Residues highlighted by black shading represent conserved amino acids, and gray shading indicates amino acids belonging to the same conservation group. SP, ER signal peptide predicted by SignalP. (**C**) Diagram shows the GLUT10/GFP/KKXX and SP/GFP/KDEL fusion proteins. Arrow indicates the putative SP cleavage site. KKXX and KDEL are the ER retention signals. (**D**) Confocal images of MOVAS cells expressing the indicated GFP fusion proteins stained with ER Tracker Red. *Green,* GFP; *magenta*, ER Tracker Red; *white,* merged. Scale bars, 10 µm. (**E**) Confocal images of MOVAS cells expressing the indicated GFP fusion proteins stained with MitoTracker Red. *Green,* GFP; *magenta*, MitoTracker Red; *white,* merged. Scale bars, 10 µm. (**F**) Percentage of GFP fusion proteins colocalized with ER in cells, as in **D**. Data represent the mean ± SEM, n = 5-10 independent experiments with 10 cells per group in each experiment. (**G**) Percentage of GFP fusion proteins colocalized with mitochondria of cells, as in **E**. Data represent the mean ± SEM, n = 7-9 independent experiments of 10 cells per group in each experiment.

To further investigate whether retained GLUT10 in ER might affect its mitochondrial trafficking, a transmembrane protein ER retention signal, di-lysine (KKXX) motif ^19^ was fused to the C-terminus of GLUT10/GFP (GLUT10/GFP/KKXX) (Fig. 3C). The GLUT10/GFP/KKXX protein was retained in ER and exhibited reduced mitochondrial trafficking under H_2_O_2_ treatment (Fig. 3D-G), suggesting that retention of GLUT10 in the ER can impair its mitochondrial targeting.

GLUT10 contains a predicted N-linked glycosylation site (Fig. 4A), and our western blotting results showed different molecular weights of GLUT10 in mitochondrial fractions (Fig. 2M-P). Since N-linked glycosylation typically occurs in the ER and Golgi ^20^, we wondered whether mitochondrial GLUT10 might be transported through the ER-Golgi pathway and undergo N-glycosylation. To test this idea, mitochondria-enriched fractions were treated with a de-glycosylating enzyme, N-glycosidase F (PNGase F), which removes N-linked glycans ^21^. This treatment reduced the molecular weight of GLUT10 in mitochondrial-enriched fractions (Fig. 4A). Since the GFP moiety is not glycosylated ^22^, these results indicate that mitochondrial GLUT10 is indeed N-glycosylated. To confirm ER to Golgi translocation is important for GLUT10 glycosylation and targeting to mitochondria, we treated cells with brefeldin A (BFA), an inhibitor of vesicle formation and transport between the ER and Golgi ^23^. BFA treatment resulted in GLUT10 accumulation in the ER (Fig. 4B) and reduced its molecular weight in both mitochondrial and ER-enriched fractions (Fig. 4C), in line with the expected loss of N-glycosylation. Of note, the lower molecular weight GLUT10 species observed in mitochondrial fractions after BFA treatment might be derived from enhanced ER-mitochondria contact ^24^, which could potentially enhance GLUT10 presence in mitochondria-enriched fractions.

**Figure 4.**
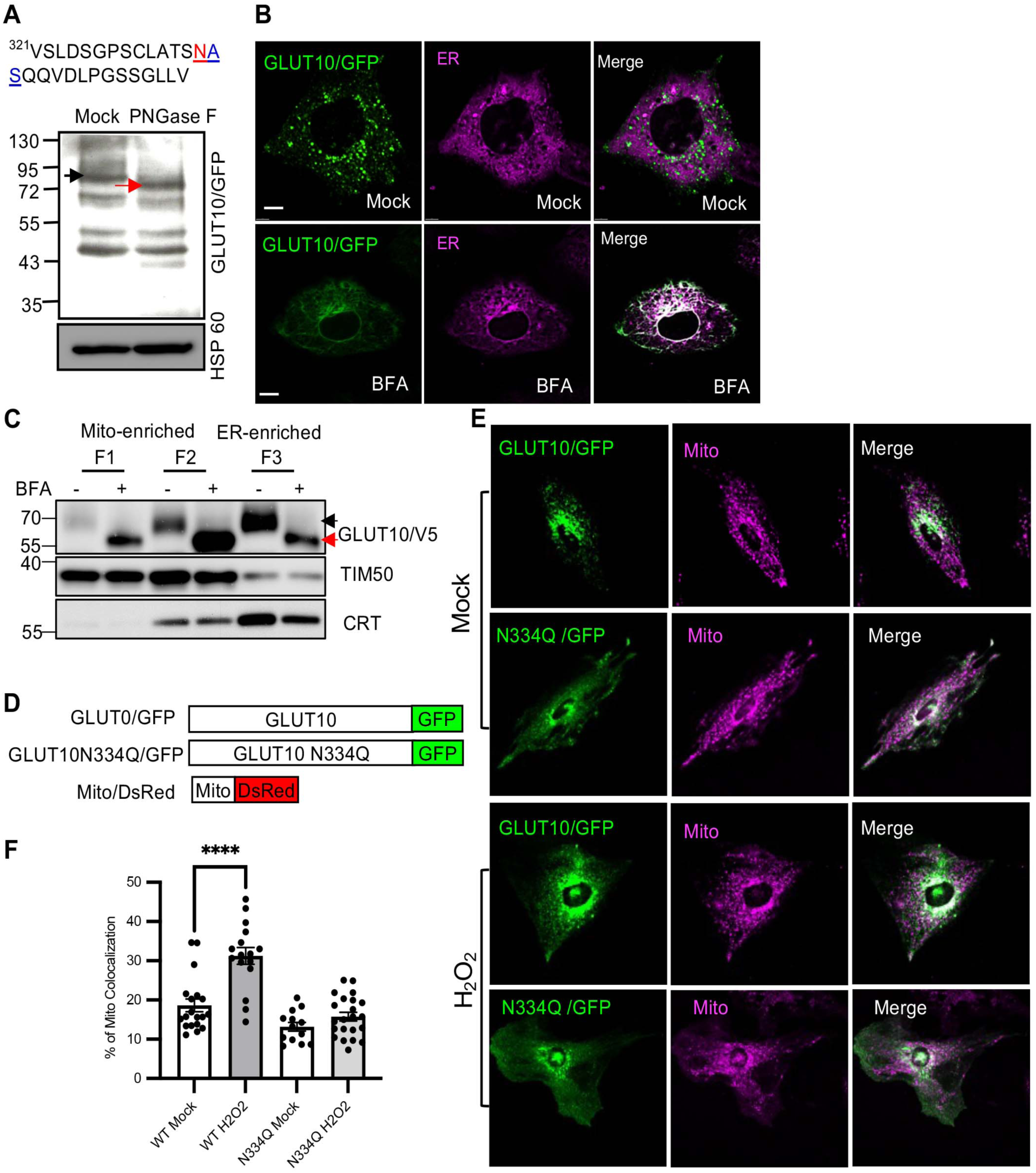
Mitochondrial GLUT10 is N-glycosylated. (**A**) The N-glycosylation site (N-X-S, underlined triplet) in mouse GLUT10 was predicted with NetNglyc1.0. The asparagine (N) predicted to be N-glycosylated is highlighted in red. Immunoblot analysis showed a molecular weight shift of GLUT10/GFP in mitochondrial-enriched fractions from GLUT10/GFP-expressing A10 cells treated with PNGaseF (indicated by black and red arrows). HSP60, Heat shock protein 60, a mitochondrial marker. (**B**) Confocal images of GLUT10/GFP colocalized with ER in A10 cells expressing GLUT10/GFP treated with or without 5 μM BFA for 1 h; ER was stained with ER Tracker Red. *Green,* GFP; *magenta*, ER Tracker Red; *white,* merged. Scale bars, 10 µm. (**C**) Immunoblots to detect molecular weight changes of GLUT10/V5 in subcellular fractions. GLUT10/V5 T-REx-293 cells were induced with TET for 12 h. After 4 h of induction, cells were treated with or without 3.56 μM BFA for 4 h. BFA was then removed, and induction continued for another 4 h. IB of V5 for GLUT10/V5; TIM50, mitochondrial marker; CRT, calreticulin, ER marker. (**D**) Diagram shows GLUT10/GFP, GLUT10 N334Q mutant fused to GFP (GLUT10 N334Q/GFP), and Mito/DsRed fusion protein. (**E**) Confocal images of A10 cells expressing GLUT10/GFP or GLUT10 N334Q/GFP and Mito/DsRed treated without (Mock) of with 100 μM H_2_O_2_ for 12 h. *Green,* GFP; *magenta*, Mito/DsRed; *white,* merged. Scale bar, 25 µm. (**F**) Quantification of the percentage of GLUT10/GFP (WT) or GLUT10 N334Q/GFP (N334Q) colocalized with mitochondria, as in **B**. Data represent the mean ± SEM, n = total of 13-22 cells from three independent experiments. Unpaired two-tailed Student’s t-test. \*\*\*\**P* < 0.0001.

Since N-glycosylation is a well-established regulator of intracellular protein trafficking ^25^, so we next tested the effect of mutating the predicted N-linked glycosylation site from asparagine (N) to glutamine (Q) in GLUT10 (GLUT10-N334Q) (Fig. 4D). The results showed that the mutation increased GLUT10 PM localization and significantly reduced its mitochondrial targeting upon H_2_O_2_ treatment (Fig. 4E and F). This suggests that GLUT10 N-glycosylation is essential for its mitochondrial targeting.

In conclusion, these findings suggest that GLUT10 is transported through the ER-Golgi system, where it undergoes N-glycosylation before being targeted to mitochondria.

### YXXΦ motif of GLUT10 mediates clathrin-dependent endocytosis from the plasma membrane (PM)

To investigate the mechanisms underlying GLUT10-containing vesicle targeting to mitochondria, we examined the potential role of an evolutionarily conserved vesicle trafficking motif (YXXΦ) identified in the C-terminal region of GLUT10 (YXXΦ, Y denotes tyrosine; X is any amino acid; and Φ is an amino acid with a bulky hydrophobic side group, such as leucine, isoleucine, methionine, valine or phenylalanine) (Fig. 5A). This motif is known to mediate rapid internalization and protein sorting from the PM to endosomal and lysosomal compartments ^26^. Deletion of the YXXΦ motif in GLUT10 (GLUT10d/GFP) (Fig. 5B) enhanced its PM localization (Fig. 5C and D). As GLUT10 is responsible for DHA uptake, and the DHA is rapidly converted to AA in the intracellular environment ^14,15^, the increased PM localization of GLUT10d/GFP was associated with increased DHA uptake and increased intracellular AA levels in MOVAS cells expressing GLUT10d/GFP (Fig. 5E). This suggests that PM localization of GLUT10 is crucial for increasing intracellular AA levels. Most importantly, the increase in GLUT10d/GFP on the PM corresponded to a decrease in mitochondrial GLUT10 targeting (Fig. 5F and G), suggesting that the deletion impairs GLUT10 endocytosis from the PM and its subsequent transport to mitochondria. The function of the YXXΦ motif in GLUT10 was further demonstrated by replacing the C-terminal 10 amino acids of GLUT1 with that of GLUT10 (GLUT1/YXXΦ/GFP) (Fig. 5H). While GLUT1/GFP was predominantly localized to PM, GLUT1/YXXΦ/GFP showed minimal PM localization and exhibited much broader intracellular distribution (Fig. 5I and J). These findings suggest that the YXXΦ motif mediates GLUT10 endocytosis from the PM, which plays a crucial role in GLUT10 mitochondrial targeting.

**Figure 5.**
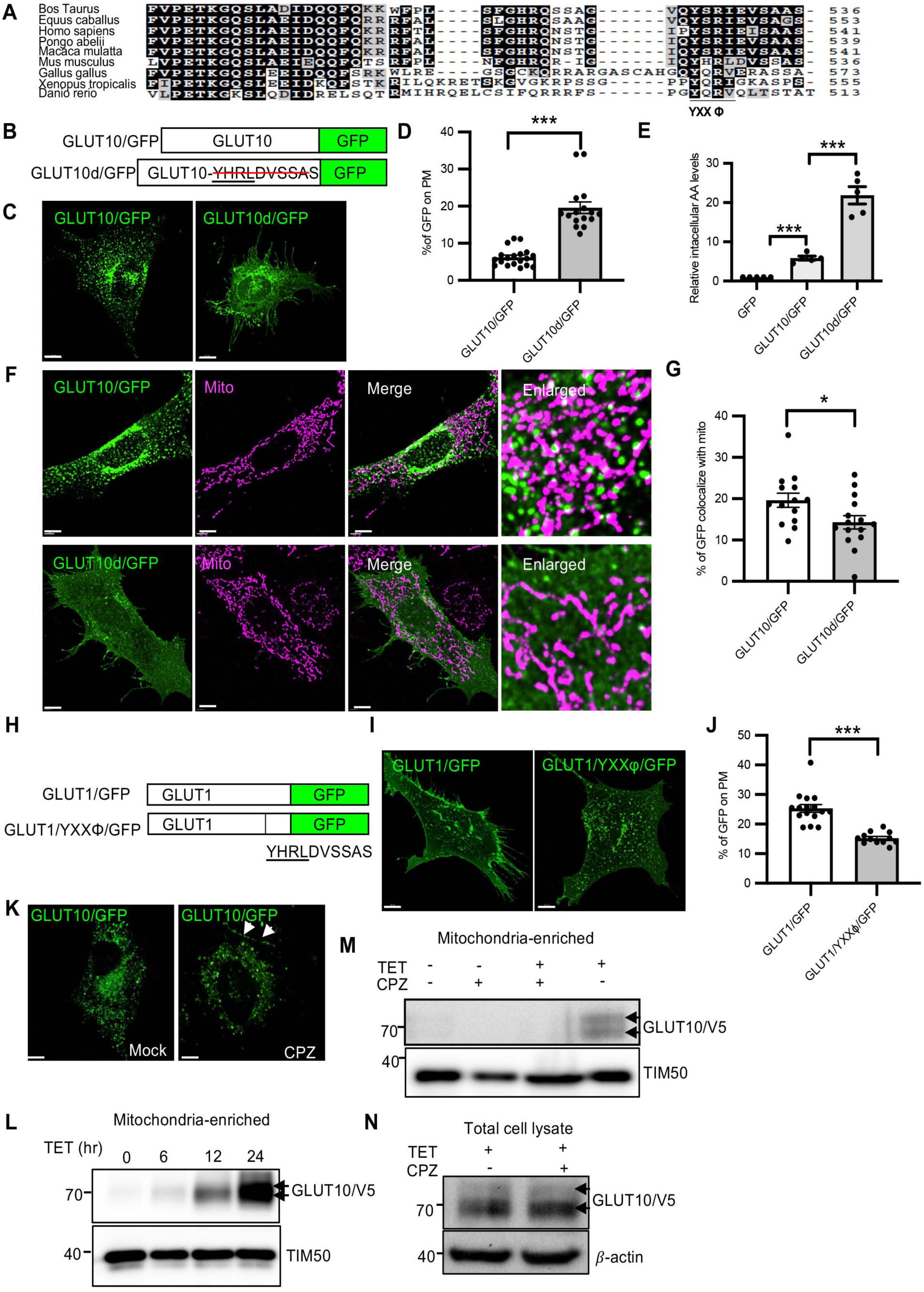
YXXΦ motif and clathrin mediate GLUT10 plasma membrane endocytosis and mitochondrial targeting. (**A**) Amino acid alignments of GLUT10 C-terminus from several species analyzed by CLUSTAL W program. The YXXΦ motif is underlined. Residues in black shading are conserved, while gray shading indicates amino acids belonging to the same conservation group. (**B**) GLUT10/GFP and YXXΦ motif-deleted GLUT10/GFP (GLUT10d/GFP) fusion proteins. (**C**) Confocal images of MOVAS cells expressing GLUT10/GFP or GLUT10d/GFP. Scale bar 10 µm. (**D**) Quantification of the relative intensity of plasma membrane-localized GFP, as in **C**. A custom ImageJ macro was used as described in Supplementary materials and Supplementary figure 8. Data are shown as mean ± SEM, n = total 16-19 cells from 3 independent experiments. (**E**) Quantification of the DHA uptake. A10 cells expressing GFP, GLUT10/GFP or GLUT10d/GFP were incubated 5 mM DHA for 30 mins, and intracellular AA levels were measured by HPLC and presented as relative levels compared to cells expressing GFP control. Data are shown as mean ± SEM from 5 independent experiments. (**F**) Confocal images of MOVAS cells expressing GLUT10/GFP or GLUT10d/GFP stained with MitoTracker Red. *Green,* GFP; *magenta*, MitoTracker Red; *white,* merged. Scale bars, 5 µm. (**G**) Quantification of the percentages of GLUT10/GFP and GLUT10d/GFP colocalized with MitoTracker, as in **F**. Data represent the mean ± SEM, n = 14-15 cells from 3 independent experiments. (**H**) GLUT1/GFP and GLUT1-YXXΦ/GFP fusion proteins. (**I**) Confocal images of MOVAS cells expressing GLUT1/GFP or GLUT1-YXXΦ/GFP. Scale bar 10 µm. (**J**) Quantification of the relative intensity of plasma membrane GFP, as in **I**. Images were analyzed using a custom ImageJ macro described in Supplementary materials and Supplementary figure 8. Data are shown as mean ± SEM, n = total 12-16 cells from 3 independent experiments. (**K**) Confocal images of GLUT10/GFP-expressing A10 cells treated with or without 20 μM CPZ for 6 h. Arrows indicate plasma membrane-localized GLUT10/GFP. Scale bar, 10 µm. (**L**) Immunoblots of GLUT10/V5 levels in mitochondria-enriched fractions from T-REx-293 cells with induced GLUT10/V5 expression at indicated time points. IB, V5 for GLUT10/V5, TIM50, mitochondrial marker. (**M** and **N**) Immunoblots of GLUT10/V5 levels in (**M**) mitochondria-enriched fractions and in (**N**) total protein lysates of T-REx-293 cells pretreated with 20 μM CPZ for 1 h before induction of GLUT10/V5 expression for 6 h. IB, V5 for GLUT10/V5, TIM50, mitochondrial marker, beta-actin served as total protein loading control. Statistical comparisons were made with two-tailed Student’s t-test in **D**, **E** and **H**. \**P* < 0.05, \*\**P* < 0.01, \*\*\**P* < 0.001.

Given that the YXXΦ motif is typically recognized by the adaptor protein-2 (AP-2) complex to facilitate clathrin-mediated endocytosis ^27^, we next tested whether GLUT10 undergoes YXXΦ-mediated PM endocytosis via a clathrin-mediated pathway. As predicted, the results showed that GLUT10/GFP colocalized with clathrin (Fig. S5A and B). Furthermore, inhibiting clathrin-AP2-mediated endocytosis with chlorpromazine (CPZ; inhibits the assembly of clathrin and AP2 on endosomal membrane ^28^) enhanced GLUT10 PM localization (Fig. 5K). These results suggest that GLUT10 undergoes PM endocytosis through a YXXΦ motif- and clathrin-mediated pathway. In addition, inhibiting clathrin-mediated PM endocytosis with CPZ prevented the accumulation of GLUT10 in the mitochondrial fraction of the T-REx-293 inducible GLUT10/V5 expression system (Fig, S6 and Fig. 5L-M). Notably, total GLUT10/V5 expression levels remained similar in both CPZ-treated and control groups (Fig. 5N). These results indicate that clathrin-mediated PM endocytosis is essential for GLUT10 targeting to mitochondria.

Taken together, our findings led us to conclude that YXXΦ motif- and clathrin-mediated PM endocytosis of GLUT10 contribute to its mitochondrial targeting. Furthermore, the PM localization of GLUT10 is crucial for enhancing DHA uptake and increasing intracellular AA levels.

### Early endosomes (EEs) mediate GLUT10 mitochondrial targeting

To further explore the mechanisms underlying targeting of endocytosed GLUT10 to mitochondria, we next isolated GLUT10-containing vesicles and conducted proteomic analyses to identify potential regulatory proteins. The proteins identified from GLUT10-containing vesicles in GLUT10/V5-expressing T-REx-293 cells included several known regulators of vesicle trafficking, such as RAB family members (RABs) (Table S2). Immunofluorescence staining showed that GLUT10 had higher colocalization with RAB5 than other RABs (Fig. S7 and 6A). We also confirmed that RAB5, a key regulator of endosome sorting and trafficking ^29^, was present in GLUT10/V5-containing vesicles from GLUT10/V5-expressing T-REx-293 cells (Fig. 6B).

**Figure 6.**
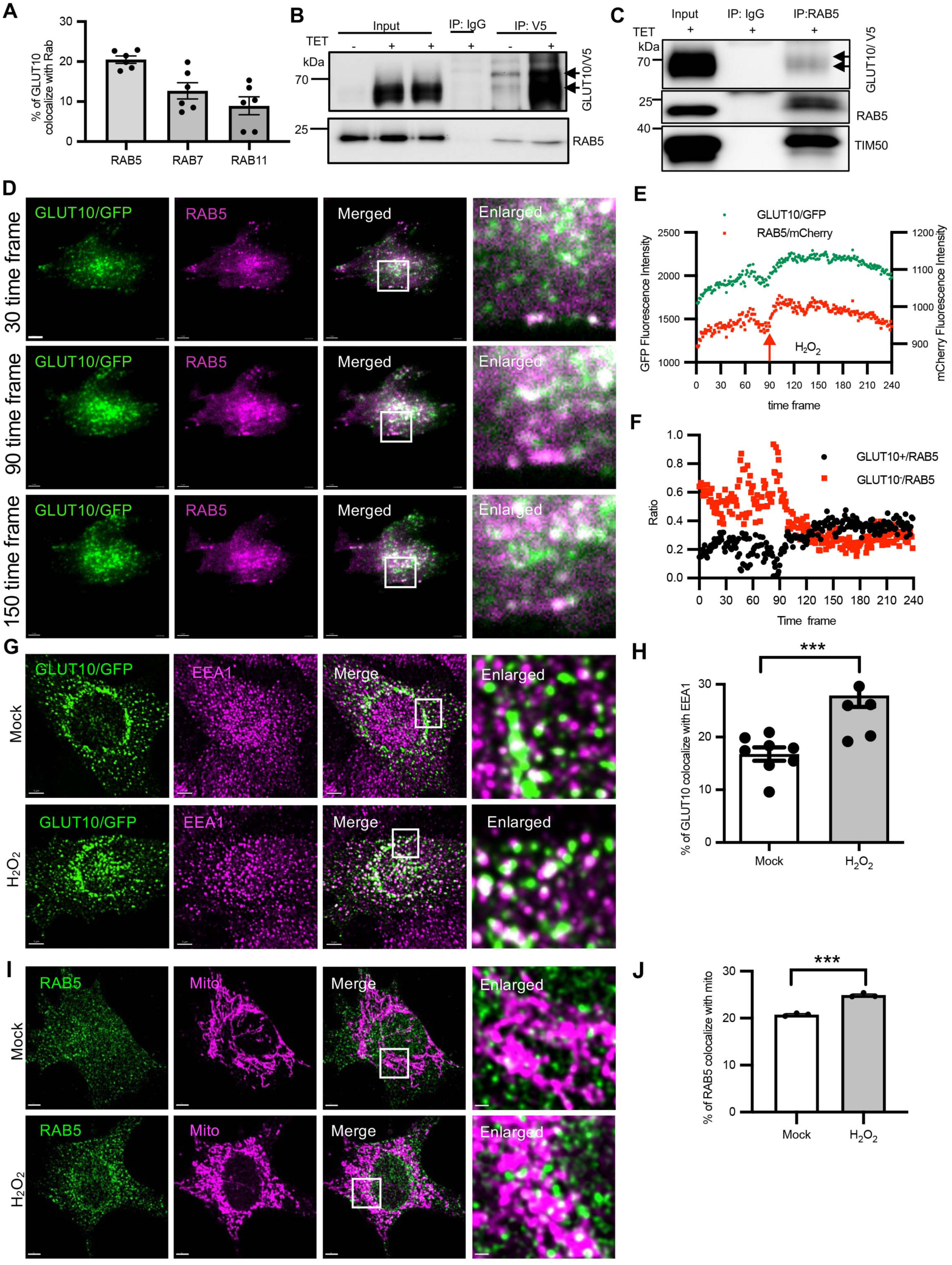
Early endosome trafficking mediates GLUT10 mitochondrial targeting. (**A**) Quantification of the percentages of GLUT10/GFP colocalized with RAB5, RAB7 and RAB11 from Supplemental figure 8. Data are shown as mean ± SEM, n = total 6 independent experiments with 10 cells per group in each experiment. (**B**) Immunoblot to analyze the presence of RAB5 and GLUT10/V5 in GLUT10-containing vesicles. GLUT10-containing vesicles were pulled down from GLUT10/V5 expressing T-REx-293 cells with V5 antibody. (**C**) Immunoblot to analyze the presence of GLUT10/V5, RAB5 and TIM50 in mitochondria-enriched fractions. RAB5 vesicles were pulled down with RAB5A antibody from mitochondria-enriched fractions. GLUT10/V5 expression was induced in T-REx-293 cells for 12 h. (**D-F**) Oxidative stress increases GLUT10 and RAB5 in plasma membrane area. (**D**) TIRF images of plasma membrane area of GLUT10/GFP (green) and RAB5A/mCherry (magenta) in GLUT10/GFP and RAB5A/mCherry expressing MOVAS cells. (**E**) Mean fluorescence intensity of GLUT10 and RAB5 in plasma membrane area before and after treatment with 50 µM of H_2_O_2_. (**F**) Ratio of colocalization of GLUT10 with RAB5 vesicles before and after H_2_O_2_ treatment. Imaging was performed at a rate of 6 frames/min (10 s/frame). Scale bar, 15 μm. (**G and H**) Oxidative stress increases GLUT10 colocalized with EEA1. (**F**) Confocal images of GLUT10/GFP colocalized with EEA1. MOVAS cells expressing GLUT10/GFP were treated with 50 µM of H_2_O_2_ for 24 h and stained by immunofluorescence for EEA1. *Green,* GFP; *magenta*, EEA1; *white,* merged. Scale bar, 5 µm. (**G**) Quantification of GLUT10 colocalized with EEA1, as in **F**. Data are shown as mean ± SEM, n = total 9-10 independent experiments with 10 cells per group in each experiment. (**I and J**) Oxidative stress increases RAB5 colocalized with mitochondria. (**H**) Confocal images of RAB5 colocalized with ATP5A1 (Mito). MOVAS cells were treated with 50 µM of H_2_O_2_ for 24 h and stained by immunofluorescence for RAB5 and ATP5A1. *Green,* RAB5; *magenta*, ATP5A1; *white,* merged. Scale bar, 5 µm. (**I**) Quantification of GLUT10 colocalized with mitochondria as in **H**. Data are shown as mean ± SEM, n = total 10 -12 independent experiments with 10 cells per group in each experiment. Statistical comparisons were made with two-tailed Student’s t-test in **G** and **I**, \*\**P* < 0.01.

We next used live imaging to monitor the trafficking of GLUT10-containing RAB5-vesicles to mitochondria. We captured image sequences showing RAB5-vesicles approaching, directly interacting and merging with mitochondria over time (Fig. S8A-D). Line-scan analysis confirmed the colocalization of RAB5-vesicles with GLUT10/GFP-containing vesicles and mitochondria (Fig. S8 E-G). Additionally, subcellular fractionation experiments in GLUT10/V5-expressing T-REx-293 cells demonstrated a gradual increase in GLUT10/V5-positive fractions colocalized with RAB5 and mitochondria-positive fractions after induction of GLUT10/V5 expression (Fig. S8H and I). Moreover, co-IP experiments confirmed direct interactions among RAB5, GLUT10/V5 and TIM50 in mitochondria-enriched fractions (Fig. 6C). Collectively, these results strongly suggest that GLUT10 is present in RAB5-vesicles, which are targeted to mitochondria.

### Oxidative stress-induced GLUT10 PM targeting, endocytosis, and endosomal trafficking to mitochondria

Since oxidative stress enhances GLUT10 mitochondrial targeting (Fig. 4D), we next examined whether oxidative stress might enhance trafficking of GLUT10, including targeting to the PM, endocytosis, colocalization with RAB5-vesicles, and mitochondria targeting. Using total internal reflection fluorescence (TIRF) microscopy, we monitored the dynamics of GLUT10 and RAB5-vesicles in the vicinity of the PM. Continuous imaging before and after H_2_O_2_ treatment at 10-s intervals revealed a rapid increase in GLUT10 signal and a slight increase in RAB5 signal upon the addition of H_2_O_2_ (Fig. 6D and E; Movie S5). These increases in signal indicated that oxidative stress enhances GLUT10 targeting to the PM and increases RAB5 localization near the PM. Furthermore, we also analyzed the colocalization of GLUT10 with RAB5 vesicles around the PM. Before H_2_O_2_ treatment, RAB5 vesicles lacking GLUT10 were abundant, as the GLUT10^−^/RAB5 ratio was high. Few RAB5 vesicles contained GLUT10, as indicated by the low GLUT10^+^/RAB5 ratio. After H_2_O_2_ treatment, the GLUT10^+^/RAB5 ratio increased, and the GLUT10^−^/RAB5 decreased (6D and F). We observed similar trends of increased GLUT10 colocalization with RAB5 vesicles after addition of H_2_O_2_ across several cells, despite variations in the intensity of GLUT10/GFP and RAB5/mCherry signals due to the transfection efficiency.

Given the fact that the H_2_O_2_ treatment increased GLUT10 endocytosis from the PM, we next analyzed GLUT10 colocalization with the early endosome marker EEA1. As expected, H_2_O_2_ treatment significantly increased colocalization of GLUT10 with EEA1, and it also increased colocalization of RAB5-vesicles with mitochondria throughout the cell (Fig. 6G-J). These results further support the idea that H_2_O_2_ treatment promotes GLUT10 PM targeting, endocytosis and colocalization with RAB5 vesicles, which targets the protein to mitochondria.

### Disrupting intracellular trafficking and subcellular localization of GLUT10 affects intracellular AA homeostasis

To further explore the physiological significance of GLUT10 intracellular trafficking, we investigated how disrupting the process affects intracellular AA homeostasis in cells cultured in AA-containing medium. Notably, 293T cells expressing GLUT10d/GFP and GLUT10N334Q/GFP, enhanced PM localization, showed significantly increased intracellular AA levels compared to control cells expressing GLUT10/GFP (Fig. 7A). These findings highlight the crucial role of GLUT10 intracellular trafficking in regulating intracellular AA homeostasis. Furthermore, these results suggest that PM localization of GLUT10 play important role in enhancing DHA uptake and elevating intracellular AA levels.

**Figure 7.**
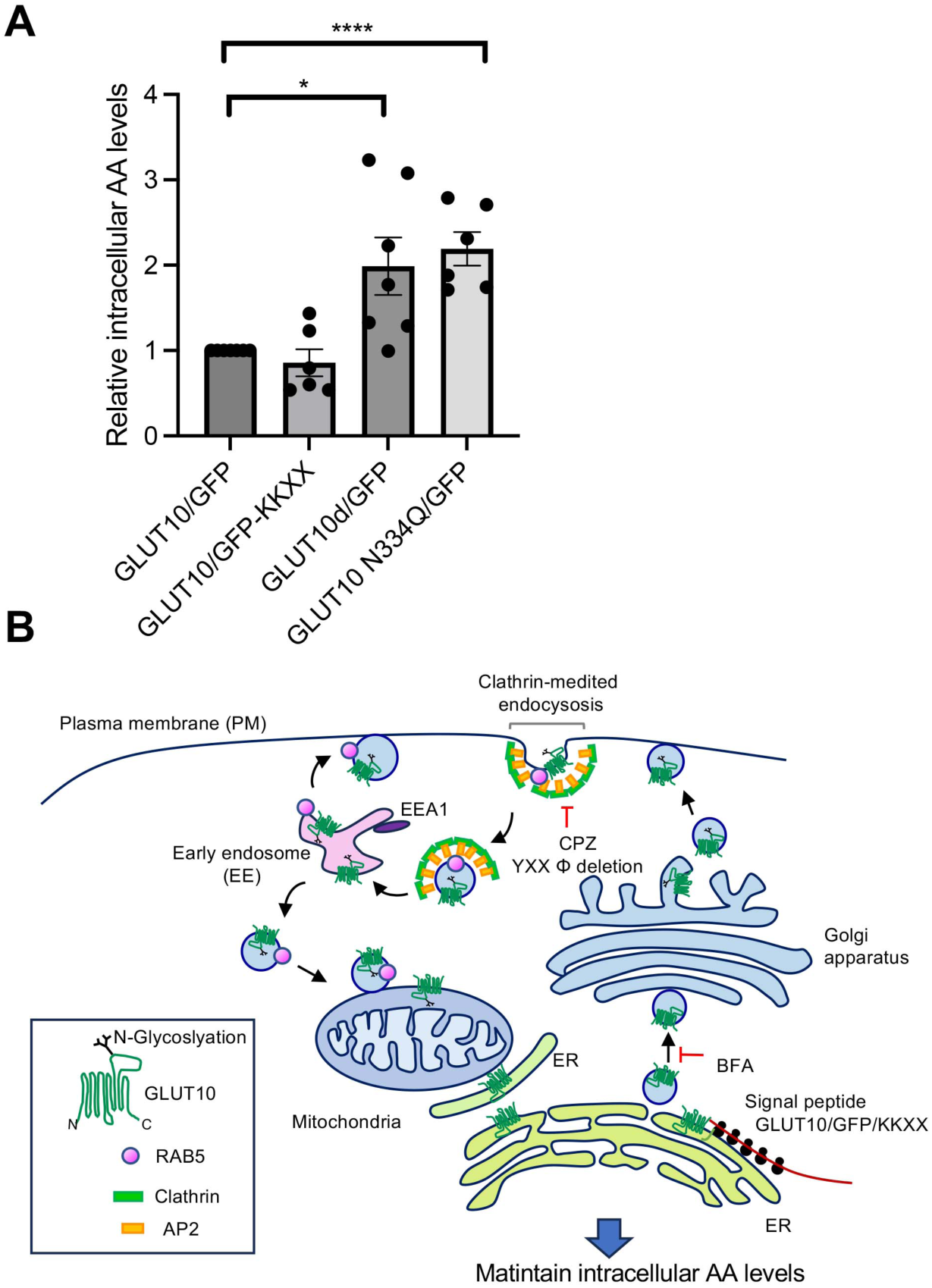
Disrupting intracellular trafficking and subcellular localization of GLUT10 affects intracellular AA levels; proposed model of GLUT10 intracellular trafficking. (**A**) AA hemostasis was analyzed by determining the intracellular AA levels in 293T cells transfected with different versions of GLUT10 and cultured in medium containing 75 µM AA for 2 days. N = 6 independent experiments for each group. Statistical comparisons were made with two-tailed Student’s t-test. \**P* < 0.05, \*\**P* < 0.01, \*\*\**P* < 0.001. (**B**) Model illustrates the transport of GLUT10 from ER to mitochondria. The SP of GLUT10 targets the protein to the ER/Golgi, where it undergoes N-glycosylation. Treatment with BFA results in retention of GLUT10 in ER-like structures. GLUT10/GFP/KKXX fusion proteins restrain GLUT10 in the ER system and reduce GLUT10 mitochondrial targeting. Deletion of YXXΦ motif in GLUT10 or inhibition of clathrin-mediated plasma membrane (PM) endocytosis by CPZ treatment both result in GLUT10 enrichment at the PM and reduced GLUT10 mitochondrial targeting. Endocytosed GLUT10 is localized to RAB5-vesicles and targeted to mitochondria. Notably, GLUT10 in the mitochondrial fraction is N-glycosylated. Overall, we propose that GLUT10 is first targeted to the ER and Golgi for N-glycosylation, then targeted to the PM, endocytosed from PM via its YXXΦ motif- and the clathrin-mediated pathway, and finally targeted to mitochondria through an endosomal-mediated pathway.

## Discussion

It is widely accepted that distinctly different mechanisms mediate the targeting of proteins to the endomembrane system and to mitochondria. In this study, we describe a unique trafficking pathway in which GLUT10 is first targeted from ER-Golgi to the PM; it is then endocytosed to early endosomes and delivered to the mitochondrial outer membrane through endosomal trafficking (Fig. 7B). Our study uncovers three key findings: (1) the interplay between the endomembrane system and mitochondria enables efficient redistribution of N-glycosylated proteins, facilitating their dynamic relocation across cellular compartments; (2) GLUT10 is newly identified at the PM, where it plays an essential role in maintaining intracellular AA homeostasis by enhancing DHA uptake; and (3) under stress conditions, this trafficking pathway is enhanced, supporting cellular adaptation to stress by modulating intracellular AA levels. Together, these findings reveal a unique mechanism for protein trafficking that links endomembrane-mitochondrial interactions to cellular homeostasis and stress resilience, advancing our understanding of GLUT10’s functional roles and providing insights with broader physiological relevance beyond ATS.

GLUT10 exhibits high expression in smooth muscle cells^14,15^, so we mainly used ASMCs (A10 and MOVAS) as a model system for our experiments. Since there is no commercially available mouse/rat-specific GLUT10 antibody, we instead utilized a GLUT10/GFP fusion protein to inspect GLUT10 subcellular localization and intracellular trafficking in MOVAS and A10 cells. To avoid discrepancies that may arise from the use of a large GFP tag or continuous protein expression, an inducible T-REx-293 system expressing GLUT10/V5 was also used as a control. Additionally, some experiments were validated by detecting endogenous GLUT10 in hASMCs and 293T cells. Although these various experimental systems consistently supported our major conclusions, we did observe some variations between cell types. For instance, the different cell types exhibited different levels of N-glycosylation modifications and mitochondrial targeting, as evidenced by differences in the molecular weight and abundance of GLUT10 in hASMCs and 293T cells (Fig. 1S). While we observed GLUT10 trafficking from the endomembrane system to mitochondria in both ASMCs and 293T cells, our findings also indicate that the subcellular distributions of GLUT10 in different cell types may be diverse due to their distinct cellular characteristics.

In our experiments, we investigated the intracellular localization of GLUT10 and its functional implications. Confocal imaging illustrated the overall distribution of GLUT10, while electron microscopy provided detailed localization patterns. Similar to GLUT4, GLUT10 undergoes intracellular redistribution to fulfill its function. GLUT4 primarily resides in the trans-Golgi network, endosomes, and storage vesicles, and is translocated to the PM in response to insulin to enhance glucose flux ^30^. GLUT10, along with GLUT6, GLUT8 and GLUT12, belongs to the class 3 GLUTs, all of which exhibit intracellular localization^15,31,32^, and their intracellular function and regulation remain largely unclear. We demonstrated that stress-induced GLUT10 located from ER to mitochondria in agreement with our previous findings ^14,15^; in addition, we observed GLUT10 at PM, in endocytosis early endosomes, and in early endosomes close contact with mitochondria, suggesting the communication between the endomembrane and mitochondria for protein redistribution. Most importantly, under stress conditions, GLUT10 targeting to the PM and early endosomes was significantly enhanced, facilitating its mitochondrial targeting. This PM localization of GLUT10 markedly increases DHA uptake, thereby boosting intracellular AA levels. We provide the first evidence for GLUT10 at the PM and highly its critical role in maintaining AA homeostasis through enhanced PM localization and mitochondrial targeting under stress.

Previous studies have identified mitochondrial N-glycosylated proteins ^33^ and shown that N-glycosylation is essential for proper mitochondrial targeting ^6^. However, how these proteins are N-glycosylated and trafficked to mitochondria remains unknown. Our study identifies a pathway by which N-glycosylated GLUT10 is trafficked from the ER-Golgi to mitochondria via the plasma membrane (PM) and endosomes, providing insight into the mitochondrial targeting of N-glycosylated transmembrane proteins. Disrupted GLUT10 N-glycosylation increased its PM retention and reduces mitochondrial targeting, suggesting that N-glycosylation of GLUT10 may facilitate both endosomal sorting and mitochondrial targeting. Interestingly, we observed two molecular forms of GLUT10 in mitochondria. Notably, under BFA treatment, only the lower molecular weight form was present (Fig. 4C), and GLUT10 was also observed at ER-mitochondria contact sites (Fig. 2F). These findings suggest that in addition to the fully N-glycosylated form, a partially glycosylated form of GLUT10 in mitochondria may reach mitochondria directly from ER-mitochondria contact sites. While the functional implications of these different GLUT10 forms in mitochondria remains to be clarified, our data show that the ER-Golgi N-glycosylation and PM targeting are functionally significant for GLUT10. These findings reveal a previously unexplored mechanism for targeting N-glycosylated transmembrane protein to mitochondria, raising questions about whether similar trafficking routes exist for other mitochondrial N-glycosylated transmembrane proteins.

The endosomal targeting of GLUT10 is characterized by the presence of the tyrosine-based sorting motif, YXXΦ, in the C-terminus of the protein, which mediates clathrin-dependent endocytosis from PM. The YXXΦ motif is known to mediate glycoprotein internalization, intracellular sorting, and targeting to lysosomes or lysosome-related organelles ^34 26^. The C-terminal tyrosine motif may make GLUT10 different from other intracellular localized GLUTs. Both N- and C-terminal internalization signals are present in GLUT4 (FQQI and di-leucine motifs) ^35^ and GLUT12 (both are di-leucine motifs) ^36^, whereas GLUT8 contains only the N-terminal di-leucine motif ^37^. Although both the tyrosine motif and di-leucine motif can mediate endocytosis from the PM, these motifs further determine the sorting of proteins to different vesicular compartments^38^. Given the localization of GLUT4 and GLUT8 to specialized vesicular compartments other than endo/lysosomes, the di-leucine motif might play a role targeting the protein to specialized compartments and also in trafficking from the trans-Golgi network to endosomes ^37,39^. Nevertheless, the tyrosine motif is mainly thought to mediate direct trafficking within endocytic/secretory pathways ^38^. The location of the tyrosine motif in transmembrane proteins, the distance from the transmembrane domain to the motif, and the number of C-terminal amino acid residues after the motif are critical determinants of specific compartment targeting (Table S3) ^26^. In GLUT10, the YXXΦ motif is uniquely positioned in the C-terminal cytosolic domain; it is followed by six residues and has 36 residues separating it from the end of the predicted transmembrane domain, in contrast with previously identified YXXΦ motifs in other proteins (Table S3). While further studies are required to understand how the tyrosine motif mediates endosomal sorting of GLUT10, our findings provide insights into its potential role in endosomal trafficking.

Recent studies have highlighted the crucial functional consequences of interactions between mitochondria and endosomes/lysosomes. Transient interactions between these organelles have been shown to play functional roles in lipid and ion transfer along with mitochondrial quality control ^40^. Furthermore, oxidative stress is known to increase the RAB5 endosome interactions with mitochondria ^41^ and precede mitochondrial outer membrane permeabilization ^42^, suggesting that RAB5 endosomes target to mitochondria and regulate cellular functions in response to environmental changes. Our data reveal that endosomes mediate GLUT10 mitochondrial trafficking, and oxidative stress enhances GLUT10 localization to early endosomes as well as its mitochondrial targeting. These findings suggest that an interplay between endosomes and mitochondria enables rapid redistribution of a transmembrane protein in response to oxidative stress. This discovery adds a new dimension to our understanding of the functional interactions between mitochondria and endo/lysosomes. However, the exact mechanisms by which GLUT10-containing RAB5 endosomes are targeted to mitochondria require further elucidation.

In summary, we provide evidence supporting the existence of a protein-targeting pathway linking the endomembrane system to mitochondria. The dynamic subcellular localization of GLUT10 in the endomembrane system, PM, and mitochondria might fine-tune GLUT10-mediated transport of DHA and titrate intracellular and compartmental AA levels in response to environmental conditions. Although the detailed mechanisms of GLUT10 targeting via endosomes-mitochondria interaction require further investigation, our study uncovers a previously unknown layer of intrinsic regulation that controls the subcellular distribution of an N-glycosylated transmembrane protein functional in different subcellular compartments.

## Acknowledgements

This study was supported by grants from Academia Sinica, Taiwan (AS 105-TP-B04), the Ministry of Science and Technology (MOST), Taiwan (MOST 108-2320-B-001-022) and National Science and Technology Council (NSTC), Taiwan (NSTC 113-2320-B-001-008) to Y.C.L. The funders had no role in study design, data collection, and analysis, decision to publish, or manuscript preparation. We thank the core facilities at the Institute of Cellular and Organismic Biology, Academia, for the technical support in various imaging and biochemistry instruments. We acknowledge Dr. Marcus Calkins for English editing.

## Author contributions

Conceptualization, Y.C.L. and A.C.J.; Methodology, Y.C.L., A.C.J., YWS., Y.F.J., P.Y.L, C.Y.F., S.C.H., W.C. H, and W. C. C.; Investigation, A.C.J., Y.W.S., H.W.L., M.Y.T., Y.F.J., S.C.H., P.Y.L, C.Y.F. W.C. H, and W. C. C.; Writing-Original Draft, Y.C.L. and A.C.J.; all authors Review & Editing the Manuscript; Funding Acquisition, Y.C.L.; Supervision, Y.C.L.

## Materials and Methods

### Cell culture and transfection

Mouse aortic smooth muscle (MOVAS) cells (CRL-2797), rat aortic smooth muscle (A10) cells (CRL-1476), and HEK293T cells (CRL-11268) were obtained from ATCC (American Type Culture Collection, ATCC, Manassas, VA, USA) and maintained in Dulbecco’s Modified Eagle Medium (DMEM, Gibco) containing 10% fetal bovine serum (FBS) (Gibco) and 1% penicillin and streptomycin (Gibco). Human aortic smooth muscle cells (hASMC, 354-05A, Cell Applications) were maintained in SMC growth medium (311-500, Cell Applications). T-REx-293 cells expressing GLUT10/V5 were generated using mouse *Slc2a10* (NM_130451) cDNA and Flp-In™ T-REx™ System (K650001, Invitrogen). A stable clone was selected with 200 μg/ml of Hygromycin B. GLUT10/V5 expression was induced by 5 μg/ml tetracycline. Cell transfections were carried out using Lipofectamine^TM^ 3000 reagent (L3000001, Invitrogen). All cells were cultured at 37°C in a humid atmosphere with 5% CO_2_ and were confirmed to be negative for mycoplasma contamination.

### Antibodies and reagents

Primary antibodies included the following: V5 (R960-25, Invitrogen), GFP (ab6556, abcam), hGLUT10 (ab110528, Abcam), TIM50 (22229-1-AP, Proteintech), TOM20 (sc-11415, Santa Cruz Biotechnology), ATP5A1/mitochondrial Complex V (459240, Invitrogen), calreticulin (06-661, Merck), KDEL (ADI-SPA-827, Enzo), beta-actin (GTX110564, GeneTex), GAPDH (10494-1-AP, Proteintech), RAB5 (ab18211, Abcam), RAB5A (11947-1-AP, Proteintech), and EEA1 (ab109110, abcam). Secondary antibodies included HRP-conjugated goat anti-mouse IgG (A0168, Sigma) and goat anti-rabbit IgG (A0545, Sigma). Fluorophore-labeled secondary antibodies (Invitrogen) included AF568-goat anti-mouse IgG (A-11004) and AF 647-goat anti-rabbit IgG (A-21245). Organelle trackers (Invitrogen) included Mitotracker (M7512), ER tracker (E34250) and Golgi tracker (D7540), as well as Alexa Flour ^TM^ 568-conjugated transferrin (T23365). The PNGase F glycan cleavage kit was purchased from Gibco (A39245). Brefeldin A (B7651-5mg), chlorpromazine (C-8138), tetracycline (T7660) and hygromycin B (10843555001) were purchased from Sigma.

### Plasmid construction

The KDEL-tagged EGFP expression vector (GFP/KDEL) was generated by amplifying full-length EGFP with a reverse primer containing a KDEL-encoding sequence. The EGFP of the pEGFP-N1 expression vector (Clontech, Mountain View, CA, USA) was then replaced with EGFP-KDEL. The GLUT10/GFP and GLUT10/GFP/KKXX expression vectors were generated by insertion of the full-length mouse GLUT10 cDNA (NM_130451.1) into the pEGFP-N1 expression vector (GLUT10/GFP) or GFP/KKXX (GLUT10/GFP/KKXX) expression vector. The SP/GFP/KDEL expression construct was generated by PCR amplification of the region encoding the first 40 amino acids of GLUT10 (containing the SP) and inserted into GFP/KDEL expression vector.

The GLUT10d/GFP construct was constructed by deleting the last 10 amino acids comprising the YXXΦ motif of GLUT10 (GLUT10d/GFP) by oligonucleotide-directed mutagenesis (QuikChange Site-Directed Mutagenesis Kit, Stratagene, Santa Clara, CA, USA). The modified gene was then inserted to pEGFP-N1 expression vector. The GLUT1/GFP construct was constructed by inserting full-length mouse *slc2a1* cDNA (NM_011400.3) into the pEGFP-N1 expression vector. The GLUT1/YXXΦ/GFP construct was created by replacing the last 10 amino acid of C-terminus of GLUT1 with the last 10 amino acids of the GLUT10 YXXΦ using two-step PCR. In Step one, the DNA fragment of *slc2a1* from 892 pb to 915pb and the DNA fragment of *slc2a10* (NM_130451.1) from 1548 bp to 1697 bp, which have overlapping sequences in the homologous region, were amplified. The two PCR products from Step one were then used as templates to generate a chimeric DNA fragment containing the 892 bp to 915 bp fragment of *slc2a1* and 1548 bp –1679 bp of *slc2a10* using the 5’ primer for *slc2a1* fragment and the 3’ primer for *slc2a10* amplification. GLUT1 cDNA of the GLUT1/GFP construct was replaced by the chimeric DNA fragment to generate the GLUT1/YXXΦ/GFP construct. The primer pairs used for GLUT10/GFP, signal peptide of GLUT10, GFP/KDEL, GLUT1/GFP, GLUT1/YXXΦ/GFP construct generation, and mutagenic oligonucleotides for creation of GLUT10-deletion mutants are shown in the Supplementary Information.

Mito/DsRed in pcDNA3.1 was a gift from Dr. Ming-Der Pern, College of Life Sciences and Medicine, National Tsing Hua University. Rab5a-pmCherryC1 (addgene, #27679) was a gift from Dr. Wan-Chen Huang, Institute of Cellular and Organismic Biology, Academia Sinica.

### Immunofluorescence staining

Cells were grown on coverslips and fixed in 4% paraformaldehyde. The cells were then washed and stained with primary antibodies and fluorophore-labeled secondary antibodies. Coverslips were mounted onto glass slides using VECTASHIELD anti-fade mounting medium (H-1000-10, Vector labs).

### Confocal microscopy

Confocal images were acquired on a Laser Scanning Microscope, LSM 880 (ZEISS, Germany) using Plan-Apochromat 63x/1.4 oil immersion objective with ZEN 2.3 SP1 (black edition) software. Green and red fluorophores were excited using the 488 nm argon laser source and 561 nm diode pump solid-state laser, respectively. For live-cell imaging, the cells were grown in 35-mm glass-bottom dishes and placed in the live-cell incubation chamber. The chamber was maintained at 37°C and 5% CO_2_ to maintain normal growth conditions during the imaging process.

### Total Internal Reflection Fluorescent (TIRF) microscopy

TIRF images were captured using the TIRFM system built on an inverted microscope (Olympus IX81) equipped with a high-sensitivity EMCCD camera (iXon3 897, Andor Technology) and a UPONAPO 60x OTIRF objective lens (NA: 1.49; Olympus) coupled with Xcellence software (Olympus). 488 and 532 nm solid lasers were used to excite GFP and mCherry fluorophores, respectively. TIRF penetration depth was set at 100 nm. To capture real-time trafficking of GLUT10/GFP and RAB5A/mCherry, live cell images were recorded at a rate of 1 frame/10 s (6 frames/min). Recordings were made for 10 min before adding H_2_O_2_ and then another 10 min after adding H_2_O_2_.

### Image analysis

The acquired images were processed with Huygens software (Scientific Volume Imaging) for deconvolution. Co-localization analysis was performed using Imaris 9.9.1 (Oxford instruments). In brief, intensity-based analysis was used to calculate the percentage of colocalization between two channels. The intensity threshold for each channel was adjusted manually to best represent the fluorescence signals. Colocalized volume was obtained as a percentage of volume A above the intensity threshold with B. The fluorescence intensity of the whole cell or the plasma membrane was quantified using Fiji ^43^ with a custom ImageJ macro (see Supplementary materials). Dynamics of GLUT10/GFP vesicles in cells were characterized by tracking the positions of individual vesicles at different time points using Imaris software. To measure the fluorescence intensity of the plasma membrane area and the colocalization of GLUT10/GFP and RAB5/mCherry in time-series TIRF images, we used TrackMate ^44^ in conjunction with the Cellpose Cyto3 model ^45^ and a custom python code (see Supplementary materials).

### Immunogold and APEX2 staining electron microscopy (EM)

Immunogold labeling of permeabilized whole-mount cells for EM was performed following established procedures ^46^. Briefly, cells were grown on sapphire discs (16770158, Leica) and cryo-fixed using a high-pressure freezer (HPM100, Leica). Then, the cells were subjected to freeze substitution in an AFS2 system (AFS2, Leica) with subsequent rehydration and post-fixing. Immunogold detection was performed using anti-GFP (ab6556, Abcam), followed by 1.4 nm NANOGOLD-gold cluster labeling (#2004, Nanoprobes) and silver amplification (#2012, Nanoprobes).

The APEX2 labeling and EM sample preparation were performed as previously reported ^47^. MOVAS cells were transfected with GLUT10/APEX2 vector or APEX2 vector control. Cells were fixed and then overlaid with a solution of DAB and H_2_O_2_. APEX2 catalyzed the oxidation of DAB to generate a locally deposited DAB polymer. After labeling, cells were stained with Osmium tetroxide (OsO_4_) and Uranyl acetate (UA) to improve sample preservation and contrast. Following resin-embedding, cells were trimmed and cut into 80-nm sections using an ultramicrotome (Leica EM UC7).

Electron micrographs were recorded using FEI Tecnai microscope (FEI Tecnai G2 F20 S-TWIN, Thermo fisher) operating at 120 kV, equipped with a 2K × 2K CCD camera (US1000, Gatan) at ICOB Biological Electron Microscopy Core Facility, Academia Sinica.

### Subcellular fractionation

Subcellular fractionation was performed following a published protocol ^48^ with minor modifications. The flow-chart and detailed methods for subcellular fractionations are presented in the Supplementary materials and Supplementary figure (S1A).

### Isolation of mitochondria-enriched fractions

Mitochondria-enriched fractions were isolated using the mitochondria isolation kit for cultured cells (89874, Thermo Scientific). In brief, approximately 2 × 10^7^cells were homogenized using the reagents provided and centrifuged at 700 ×*g* at 4°C for 10 min. The supernatant was further centrifuged at 12,000 ×*g*, 4°C for 15 min to obtain the mitochondria-enriched fraction. Evidence for the enrichment of the mitochondrial fractions is shown in Supplementary figure S6.

### Western blots

Total protein lysates or organelle fractions were separated on SDS-PAGE and transferred onto a polyvinylidene difluoride (PVDF) blotting membrane (10600023, Amersham^TM^, Cytiva). The blots were detected with primary and secondary antibodies as listed in the antibodies section. Then, signals were generated with an ECL detection kit (RPN2232, Amersham^TM^).

### Proteinase K and digitonin treatment

Proteinase K digestion was performed according to the published protocol with minor modifications. ^18^. Mitochondria-enriched fractions isolated from cells were resuspended in mitochondria suspension buffer (10 mM Tris-base pH 7.4, 320 mM sucrose, 1 mM EDTA). Twenty milligrams of mitochondria in 200 µl of TD buffer (49.99 mM Tris base, 274.13 mM NaCl, 20.12 mM KCl, 13.95 mM Na_2_HPO_4_) were incubated with 1 µg/ml of proteinase K (Thermo Fisher) for 1 h at room temperature (RT) (22-27°C). The digestion was stopped by adding 2 µl of 100 mM PMSF. For digitonin treatment, mitochondria fractions isolated from cells were resuspended in 200 µl of digitonin solution (7 mg/ml) and incubated at RT with constant mixing for 1 h. Proteins were separated by SDS-PAGE and subjected to western blot analysis.

### Measurement of ascorbic acid (AA) accumulation

For DHA uptake, intracellular AA levels were measured by high-performance liquid chromatography (HPLC) as described previously ^14^. Cells were incubated in low glucose DMEM (5 mM) without FBS and containing 5 mM DHA at 37°C for 30 min. Total cell lysates were used for AA measurement. For intracellular AA homeostasis assay, AA levels were determined with an Ascorbic acid assay kit (ab65656, Abcam) according to the manufacturer’s instructions. Cells were grown in culture medium containing 75 μM AA for 48 h; AA levels were measured from whole cell lysates.

### Statistical analysis

Statistical significance between groups was analyzed by two-tailed unpaired Student’s t-test in GraphPad Prism 10 (GraphPad Software, La Jolla, CA, USA). A *p*-value less than 0.05 was considered statistically significant.

