## Supplemental data for "Plasma membrane mediated GLUT10 mitochondrial targeting regulates intracellular ascorbic acid homeostasis"

### Supplementary Materials

#### Subcellular fractionation

Subcellular fractionation was performed according to a published protocol with minor modifications [48], as shown in Supplementary figure 1A. In brief, approximately $1 \times 10^8$ cells were harvested and suspended in 2 ml of sucrose homogenization medium. Cells were lysed by homogenization using a pre-cooled Dounce homogenizer. Homogenate was centrifuged at 600 $\times g$ for 10 min, and the centrifugation was repeated until no pellet was visible. The supernatant was then further centrifuged three times at 10,000 $\times g$ at 4°C for 10 min to obtain a pellet. The pellet was further separated on 30% Percoll by centrifugation at 95,000 $\times g$ to obtain F1 and F2. The supernatant from the previous step was further centrifuged at 21,000 $\times g$ for 10 min, three times to obtain a pellet (F3); then, the supernatant was centrifuged at 200,000 $\times g$ for 60 min at 4°C to obtain a pellet (F4). The fractions were resuspended in SDS sample buffer, and organelle enrichment was analyzed by immunoblotting with specific organelle markers.

#### Immunoprecipitation and protein identification

GLUT10-interacting proteins were co-immunoprecipitated using V5 antibody (R960-25, Invitrogen) in GLUT10/V5 T-REx-293 cells expressing GLUT10/V5 after induction with 5 µg/ml tetracycline for 24 h. Cells were lysed in lysis buffer (300 mM NaCl, 50 mM Tris pH 7.5, 5% glycerol, 1% NP-40, and 1× protease inhibitor). The lysates were pre-cleaned with Protein G Mag Sepharose Xtra (GE Healthcare), and IP was performed with Protein G Mag Sepharose-conjugated V5 antibody. For the IP of GLUT10-containing vesicles, GLUT10/V5 expressing T-REx-293 cells were subjected to subcellular fractionation to collect the vesicle-enriched fraction. After pre-cleaning with Protein G Mag Sepharose Xtra (GE Healthcare), the GLUT10-containing vesicles were pulled down with Protein G Mag Sepharose-conjugated V5 antibody. The GLUT10-interacting proteins and proteins from the GLUT10-containing vesicles were further subjected to MS analysis by Orbitrap Fusion Lumos in the Proteomics Mass Spectrometry Common Facility, Institute of Biological Chemistry, Academia Sinica. Also, mitochondria-enriched fractions from GLUT10/V5 expressing T-REx-293 cells were subjected to IP using RAB5 antibody to pull down RAB5-associated mitochondria and analyzed for V5 and TIM50 by immunoblotting.

#### Determining the GFP intensity at the plasma membrane

Image analysis was performed using FIJI [43] with a custom ImageJ macro (Source code: https://github.com/WeiChenChu/Glu10_Membrane_Signal_Quantification). Briefly, a z-projection using the 'Sum Slices' method was applied to the GFP channel images. A duplicate of the projection image was created and processed to generate a binary mask using Gaussian blur and Huang thresholding [49]. The mask was further refined by applying the 'fill-hole' function and performing size filtering using the 'analyze particles' function, which resulted in a whole cell mask. The cytosol mask was then derived by iteratively applying the 'Erode' operation to the whole cell mask for 10 cycles. The cell membrane mask was subsequently created by subtracting the cytosol mask from the whole cell mask. Selections for measurements were established for specific regions within the resultant masks and saved as Regions of Interest (ROIs) in FIJI. GFP intensity measurements were conducted on the original projection image, based on the ROI list.

#### Determining Plasma membrane fluorescence intensity in time series TIRF images

Briefly, the RAB5 channel image was pre-processed with a median filter (radius = 4) to reduce noise. The filtered image was then processed using TrackMate with the Cellpose

Cyto3 model to segment the plasma membrane region. The resulting label image was converted into an ROI list corresponding to each time frame, using the 'Labels to 2D ROI Manager' function in the BioVoxxel 3D Box plugin (https://biovoxxel.github.io/bv3dbox/). Signal intensity within each ROI at each time point was measured using the FIJI ROI Manager.

**Determining colocalization in time series TIRF images**

TIRF images of GLUT10/GFP and RAB5A/mCherry were analyzed to assess vesicle colocalization over time. Initially, a cell mask was generated from the RAB5A/mCherry images using TrackMate-Cellpose (Cyto3 model) after applying a median filter with a radius of 4 pixels; this mask defined ROIs and was exported as a time-series mask image. In cases where the Cellpose analysis was not working well but the cell maintained a consistent shape across different time points, a manually defined mask image was used. The GLUT10/GFP, RAB5A/mCherry, and cell mask images were processed using custom Python code (available from the GitHub repository: https://github.com/WeiChenChu/TIRF_vesicle_colocalize_analysis) for colocalization analysis. Briefly, for each time point, Glut10-GFP images were processed with a Difference of Gaussian (DoG) filter to reduce noise and background while enhancing vesicular structures. The DoG-filtered Glut10 images were multiplied by the cell mask to isolate intracellular regions and subsequently segmented using Voronoi Otsu Labeling. RAB5A/mCherry images underwent similar DoG filtering and cell mask application, followed by thresholding using the triangle method to generate binary RAB5 masks. Colocalization was assessed by calculating the mean intensity overlap between GLUT10 labels and Rab5 masks (as overlap score, "1" means the Glut10 label is full overlap with the Rab5 mask); vesicles with an overlap score between 0.5 and 1.0 were classified as colocalized. Those between 0.01 and 0.5 were classified as partially colocalized, and those with zero overlap were classified as non-colocalized. The numbers of vesicles in each category were quantified across time points, and a colocalization ratio (colocalized vesicles divided by total Glut10 vesicles) was calculated. Data were exported to Excel files for further analysis.

**Generation of RAB5/CFP construct**

The RAB5/CFP expression vector was generated by inserting full-length rat Rab5 cDNA (AF027935.1) into the pECFP-C1 expression vector (Clontech, USA).

**Oligonucleotide primer pairs**

***Primers for plasmid construction***

Mouse *slc2a10* amplification for GLUT10/GFP construction:

Forward, 5'-CTC GAG ATG GGC CTT CGC CCA GCT GTC CT-3'

Reverse, 5'-GGA TCC GAG GAG GCT GAG GAG ACA TC-3'

Signal peptide amplification for SP/GFP/KDEL construction:

Forward, 5'-GAT CTC GAG ATG GGC CTT CGC CCA GCT GTC CT-3'

Reverse, 5'-GCA AGG ATC CCC GAA GTT CAG CTG GAG TGG-3'

KDEL-tagged GFP:

Forward, 5'-CGG GAT CCA CCG GTC GCC ACC-3'

Reverse, 5'-GCT GAT TAT GAT GCG GCC GCT TAG AGT TCA TCC TTG TAC AGC TC-3'

Mouse *slc2a1* amplification for GLUT1/GFP construction:

Forward, 5’-CTCGAG ATGGATCCCAGCAGCAAGAAG-3’

Reverse, 5’-GAATTC G CACTTGGGAGTCCGCCCC-3’

Mouse *slc2a1* fragment 892 bp to 915 bp amplification:

Forward, 5’-CTGAAG AAGCTT CGAGGGACAGCC-3’

Reverse, 5’-TTTGGTTTCAGGAAC TTTGAAGTAGG-3’

Mouse *slc2a10* fragment 1548 bp to 1697 bp amplification:

Forward, 5’-TTCACCTACTTCAAA GTTCCTGAAACCAAAGGACAG-3’

Reverse, 5’-GC GAATTC G GGA GGC TGA GGA GAC ATC CAG-3’
Rat Rab5 cDNA amplification for Rab5/CFP construct
Forward: 5’-CCGGAATTCCATGGCTAATCGAGGAG-3’

Reverse: 5’-CGCGGATCCTTAGTTACTACAACACT-3’

***Primers for site-directed mutagenesis***

GLUT10d/GFP construction:

Forward, 5’-CGC ATC GGT ATT CAG TCG GAT CCA CCG GTC-3’

Reverse, 5’-GAC CGG TGG ATC CGA CTG AAT ACC GAT GCG-3’

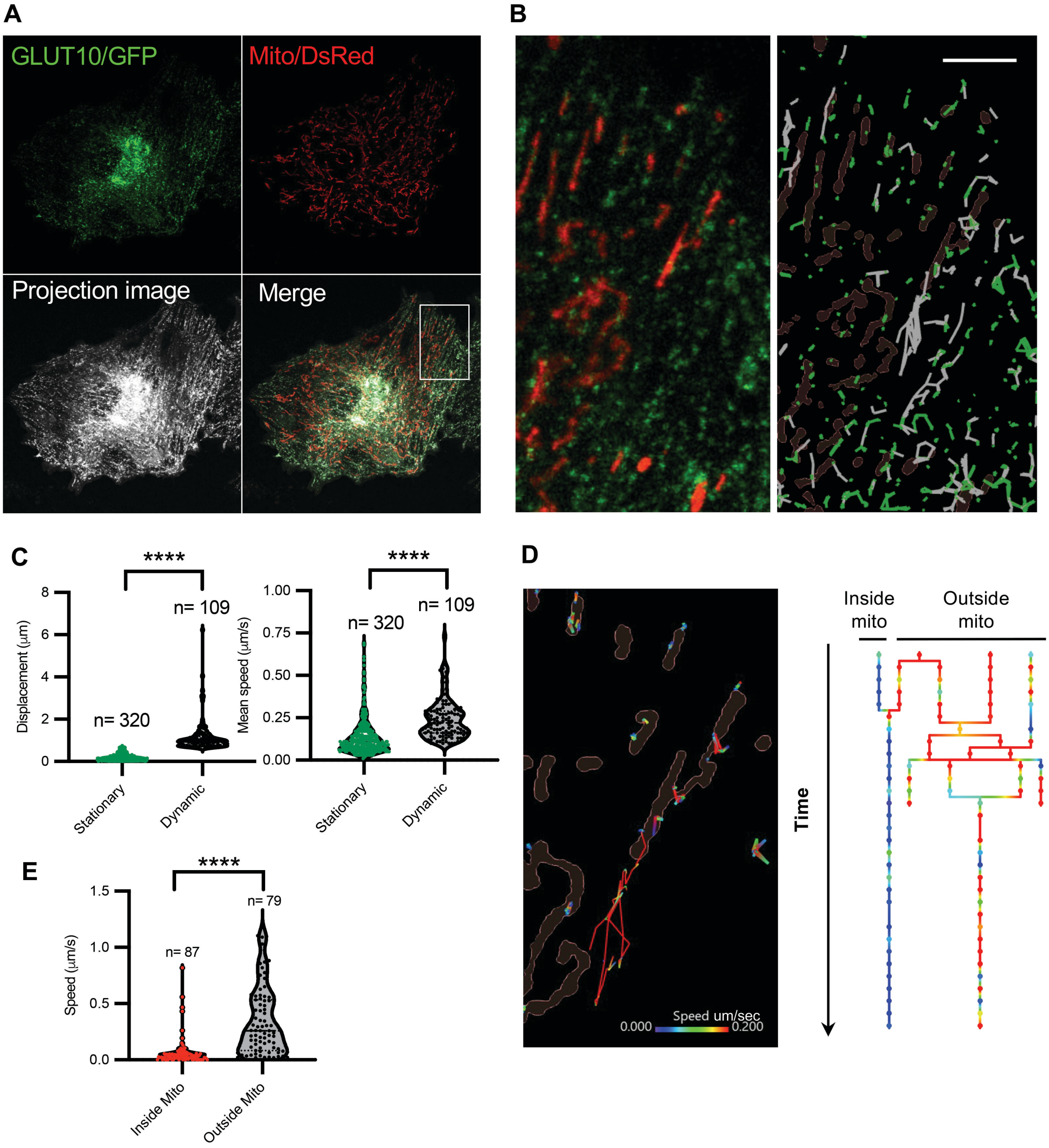


**Supplementary Figure 1. Kinetics of interaction between GLUT10-containing vesicles and mitochondria in response to $H_2O_2$ stimulation.** (**A**) Time series of confocal images shows GLUT10-containing vesicle trafficking in live A10 cells expressing both GLUT10/GFP and Mito/DsRed; cells were treated with 100 μM $H_2O_2$ and imaged at 1-s intervals for 30 s (Upper panels). The projection image was created from 10 sequential frames and shows the trafficking trajectories of GLUT10 vesicles (Lower panels). (**B**) Stationary (green) and dynamic (gray) GLUT10-vesicle trajectories were identified in the rectangle shown in **A**. A total of 429 trajectories of GLUT10 vesicles were analyzed. Scale bar: 5 μm. (**C**) The track displacement lengths and mean speeds over 30 s for stationary (green) and dynamic (white) trajectories of GLUT10-containing vesicles, as in **B**. Track displacement less than 0.67 μm over 30 s was defined as stationary. Track displacement longer than 0.67 μm over 30 s was defined as dynamic. (**D**) Representative trajectories of GLUT10-containing vesicles show the reduction of instantaneous speed after entering mitochondria. The speed of GLUT10-containing vesicle trafficking was color coded. (**E**) The mean speeds of GLUT10-containing vesicle trafficking inside and outside mitochondria. A total of 166 GLUT10-containing vesicle trajectories were analyzed, as in **A**. In **C** and **E**, data represent the mean ± SEM, n = trajectories analyzed in each group, as indicated in the figure. Statistical comparisons were made by two-tailed Student's t-test, ****$P < 0.0001$.

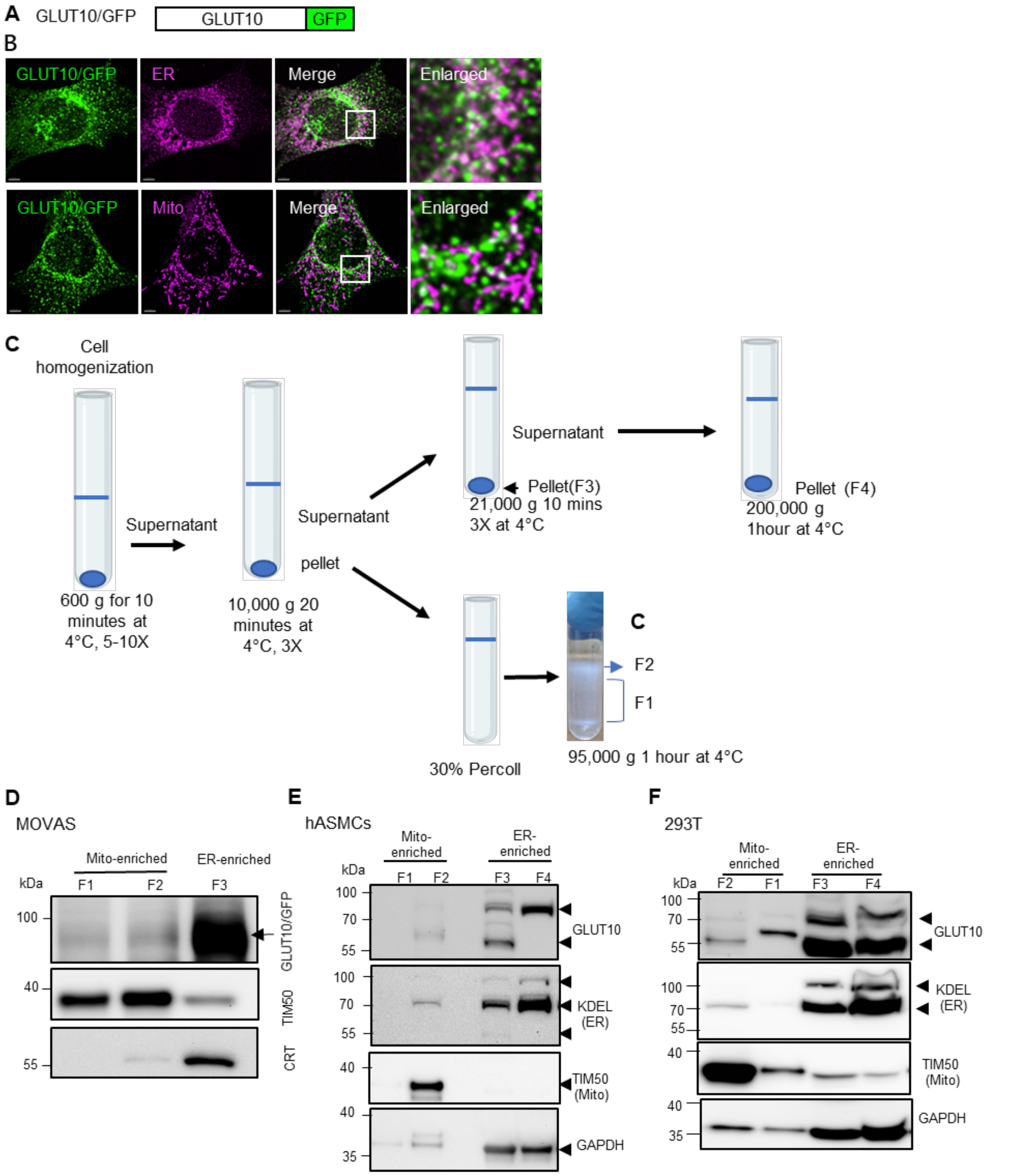


**Supplementary Figure 2. Analysis of GLUT10 subcellular distribution.** (**A**) GLUT10/GFP fusion protein. (**B**) Confocal images of GLUT10/GFP colocalized with immunofluorescence staining (IF) of subcellular compartment markers, including calreticulin (ER) and mitochondrial complex V (ATP synthase, ATP5A1) (Mito) in MOVAS cells expressing GLUT10/GFP. *Green,* GLUT10/GFP; *magenta*, IF for compartment marker; *white,* merged. Scale bars, 5 µm. **(C)** Flow-chart for subcellular fractionations. (**D**) Immunoblots of

GLUT10/GFP levels in different subcellular fractions as described in **C**; samples were derived from GLUT10/GFP-expressing MOVAS cells. TIM50, mitochondrial marker; calreticulin (CRT), ER marker. (**E** and **F**) Immunoblots of endogenous GLUT10 levels in subcellular fractions of (**E**) hASMCs and (**F**) 293T cells. KDEL, ER marker; TIM50, mitochondrial marker; GAPDH, cytoplasm marker.

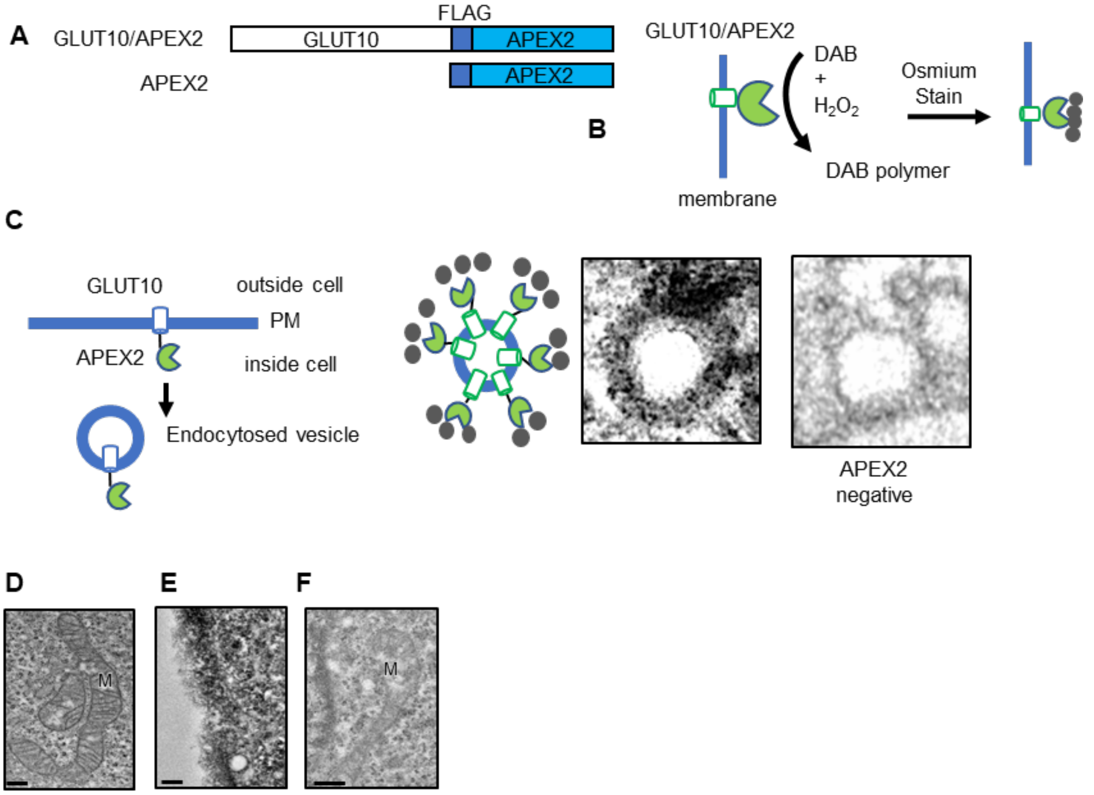


**Supplementary Figure 3. Electron microscopy (EM) characterization of GLUT10/APEX2 subcellular localization.** (**A**) Schematic of constructs expressing GLUT10/APEX2 and APEX2 control. (**B**) APEX2 catalyzes diaminobenzidine (DAB) polymerization in the presence of $H_2O_2$. The DAB polymer reacts with osmium to provide contrast for EM, indicating the localization of GLUT10/APEX2. (**C**) The schematic diagram depicts endocytosed GLUT10/APEX2-positive vesicles of enlarged EM images of GLUT10/APEX2-positive and negative endocytosed vesicles from Fig. 2I. EM image of MOVAS cells with (**D**) standard osmium fixation showing mitochondrial morphology, (**E**) APEX2 only control stained with DAB, and (F) negative control, without DAB staining. Scale bars, 100 nm.

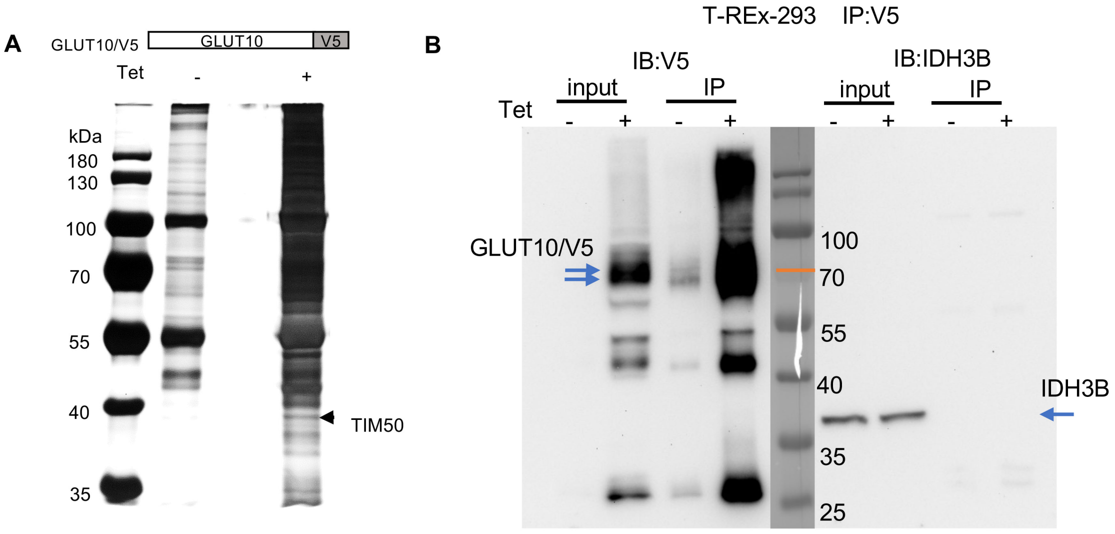


**Supplementary Figure 4. Isolation of GLUT10-interacting proteins.** (**A**) SDS-PAGE analysis of GLUT10/V5-interacting proteins pulled down with V5 from T-REx-293 cells with or without induction of GLUT10/V5 expression. Proteins were stained with Coomassie Blue. Arrow indicates TIM50, which was identified by MS analysis. (**B**) GLUT10 /V5 does not directly interact with IDH3B. Immunoblots detecting GLUT10/V5 and IDH3B protein levels in cell lysates from GLUT10/V5 expressing T-REx-293 cells before IP (Input) and after IP with V5.

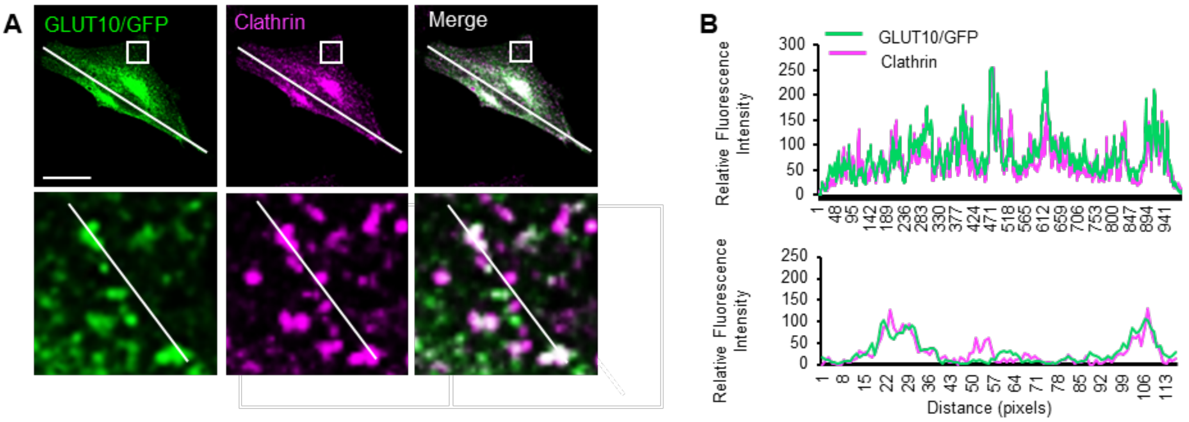


**Supplementary Figure 5. GLUT10 colocalizes with clathrin.** (**A**) Confocal images of GLUT10/GFP colocalized with immunofluorescence (IF)-labeled clathrin in GLUT10/GFP-expressing A10 cells. *Green,* GLUT10/GFP; *red*, clathrin; *yellow,* merged. Scale bar, 25 µm. (**B**) Intensity plots of line-scan analysis from **(A)** demonstrate the colocalization of GLUT10/GFP and clathrin.

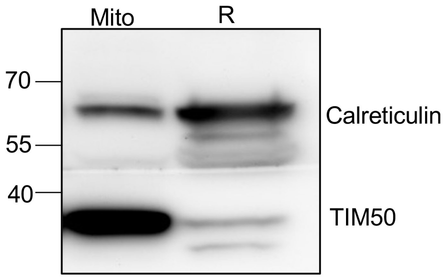


**Supplementary Figure 6. Efficiency of mitochondria isolation kit.** Immunoblots of TIM50 and calreticulin protein levels in mitochondria-enriched (Mito) and remaining (R) fractions isolated from GLUT10/V5 expressing T-REx-293 cells using the mitochondria isolation kit. Calreticulin, ER marker; TIM50, mitochondrial marker.

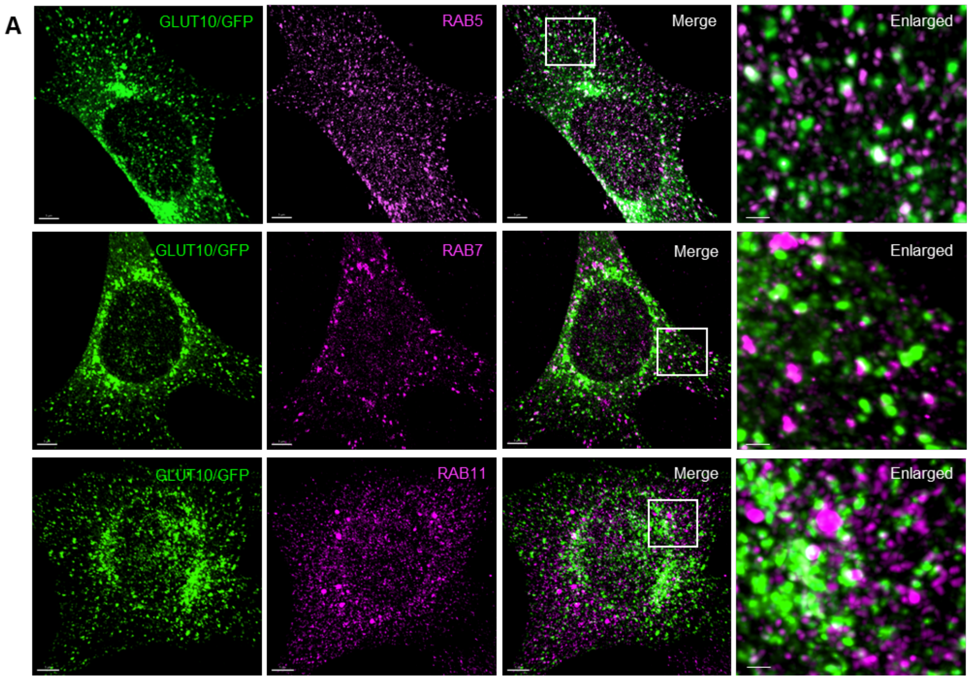


**Supplementary Figure 7. GLUT10 colocalizes with RAB5, RAB7 and RAB11.** Confocal images showing colocalization of GLUT10/GFP with RAB5, RAB7 or RAB11 in MOVAS cells expressing GLUT10/GFP and IF stained for RAB5, RAB7 or RAB11. *Green,* GFP; *magenta*, RAB5, RAB7 or RAB11; *white,* merged. Scale bar, 5 µm.

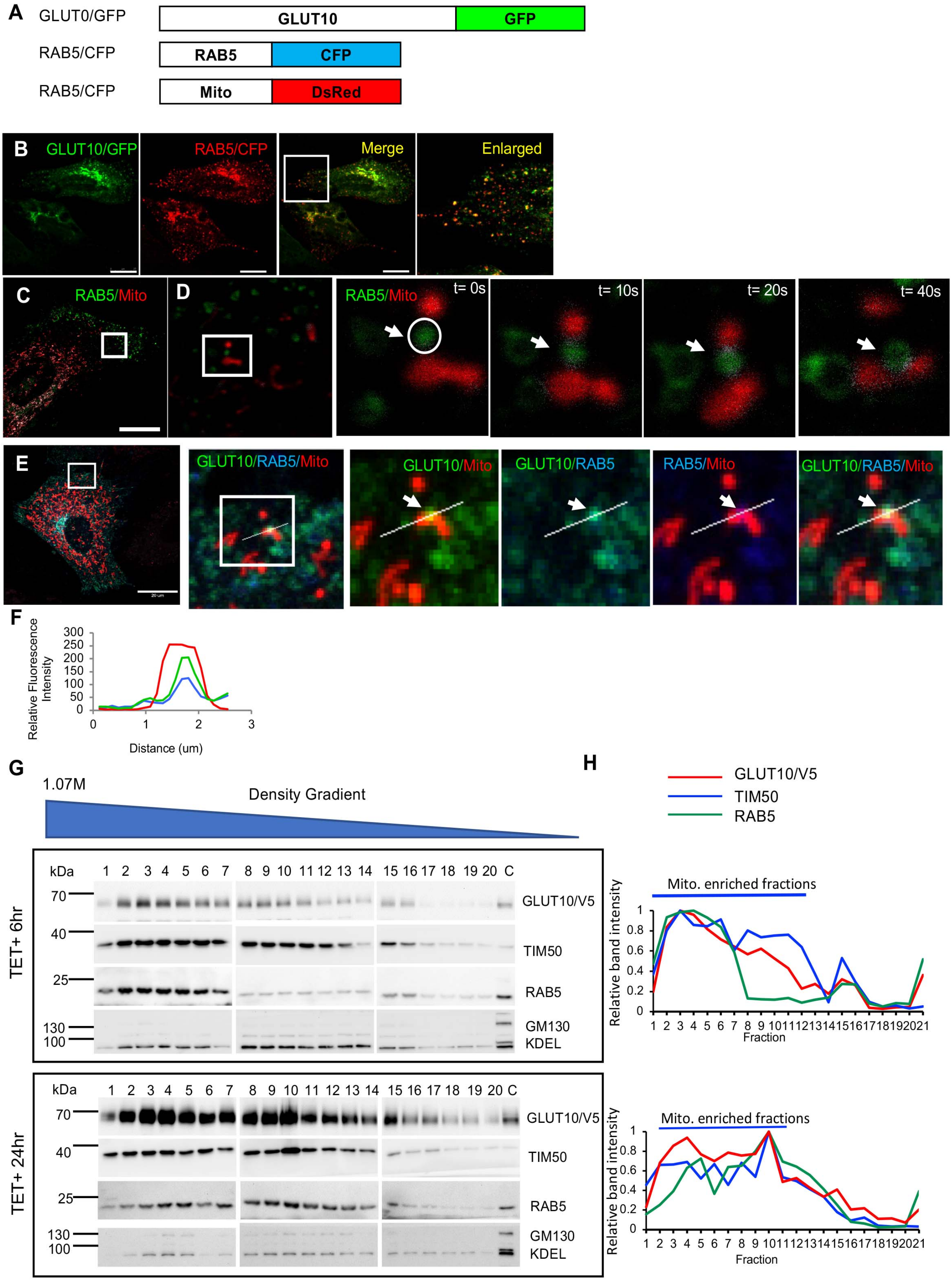


**Supplementary Figure 8. Colocalization of GLUT10 with RAB5 and mitochondria.** (**A**) Schematic representation of expression constructs for GLUT10/GFP, RAB5/CFP and Mito/DsRed fusion proteins. (**B-E**) Confocal images of live A10 cells expressing

GLUT10/GFP, RAB5/CFP, and Mito/DsRed. (**B**) Confocal images showing colocalization of GLUT10 with RAB5. *Green,* GLUT10/GFP; *red*, RAB5/CFP; *yellow,* merged. Scale bar, 10 µm. (**C** and **D**) Confocal images tracking RAB5/CFP and Mito/DsRed. *Green,* RAB5/CFP; *red*, Mito/DsRed. Scale bar, 20 µm. (**D**) Magnified time-lapse confocal images from **C** tracking the targeting of RAB5/CFP to mitochondria in live A10 cells treated with 100 µM $H_2O_2$, imaged every 10 s from t = 0-40 s. *Green,* RAB5/CFP; *red*, Mito/DsRed; *white,* merged. (**E**) Confocal images and line-scan analyses showing colocalization of GLUT10, RAB5 and mitochondria. *Green,* GLUT10/GFP; *blue,* RAB5/CFP; *red*, Mito/DsRed. Scale bar, 20 µm. (**F**) Intensity plots from the line-scan analyses of the magnified images in **E** demonstrating colocalization of GLUT10/GFP, RAB5/CFP and Mito/DsRed. (**G**) Immunoblots detecting protein levels of GLUT10/V5, TIM50, RAB5, GM130, and KDEL in subcellular organelle fractions (separated on a Percoll density gradient from GLUT10/V5 expressing T-REx-293 cells). Organelle markers: RAB5 for RAB5-positive vesicles; TIM50 for mitochondria; GM130 for Golgi, and KDEL for ER. (**H**) Quantification of the relative intensities of GLUT10/V5, TIM50, and RAB5 in different subcellular organelle fractions from **G**.

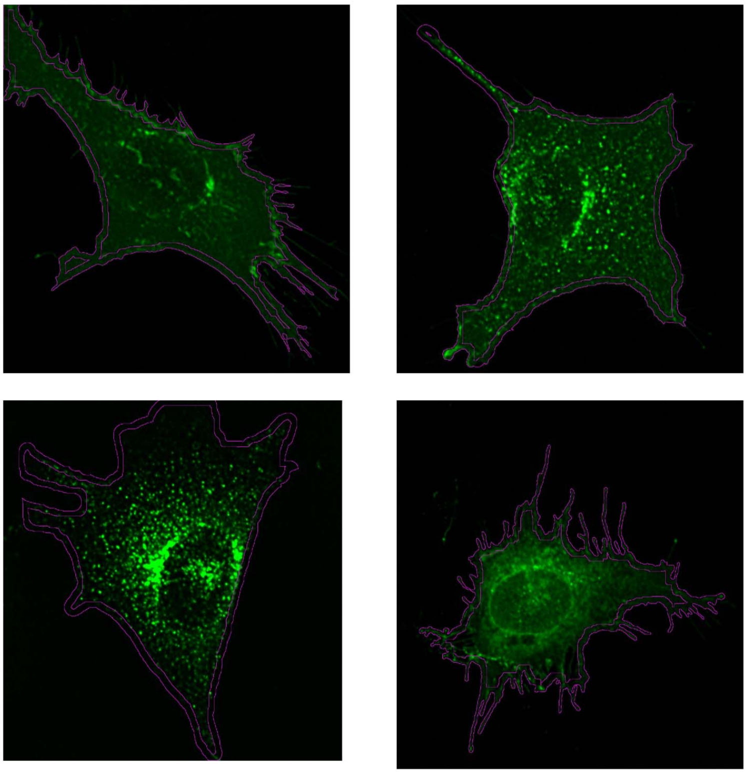

**Supplementary Figure 9. Representative images of defined plasma membrane compartment for GFP intensity calculation.** Outer border defines the whole cell. Inner border defines the cytosol. Space between the outer and inner border defines the plasma membrane of the cell. The borders were generated using a custom ImageJ macro, as described in Supplementary materials.

**Movie S1. Live time-lapse confocal imaging illustrating GLUT10/GFP targeting to mitochondria.** A10 cells co-expressing both GLUT10/GFP and Mito/DsRed. Cells were treated with 100 µM $H_2O_2$ and imaging was performed every 1 h for 7 h. Scale bar, 15 µm.

**Movie S2. Live time-lapse confocal imaging illustrating GLUT10/GFP-containing vesicles interacting with mitochondria.** A10 cells expressing both GLUT10/GFP and Mito/DsRed, Cells were treated with 100 μM $H_2O_2$; imaging was performed every 1 s for 171 s. Scale bar, 15 μm.

**Movie S3. Enlarged live time-lapse confocal imaging illustrating GLUT10/GFP-containing vesicles approaching, making direct contact with, and merging with mitochondria.** A10 cells expressing both GLUT10/GFP and Mito/DsRed, Cells were treated with 100 μM $H_2O_2$; imaging was performed every 1 s for 171 s. Scale bar, 5 μm.

**Movie S4. Electron tomogram reconstruction of MOVAS cells expressing GLUT10/APEX2, illustrating the endocytosed vesicles.** Cells were stained with DAB. EM images show strong EM contrast signals indicating the localization of GLUT10/APEX2.

**Movie S5. Live time-lapse TIRF microscopy imaging illustrating GLUT10/GFP in the plasma membrane region.** MOVAS cells co-expressing both GLUT10/GFP and RAB5A/mCherry. An initial 15 min recording shows control cell without $H_2O_2$ addition. Then, 50μM $H_2O_2$ was added, and recording was continued for another 25 min. Imaging was performed at the rate of 6 frames/min (10 s/frame). Scale bar, 15 μm.

**Table S1: Mitochondrial proteins interacting with GLUT10.** GLUT10-interacting proteins were identified and annotated as mitochondrial proteins through Gene Ontology (GO) annotation.

| No. | Gene | Protein | Sub-mitochondrial localization |
|---|---|---|---|
| 1. | AARS1 | Alanine--tRNA ligase, cytoplasmic | ND |
| 2. | AARS2 | Alanine--tRNA ligase, mitochondrial | ND |
| 3. | ABCB7 | Iron-sulfur clusters transporter ABCB7, mitochondrial | MIM |
| 4. | ABCD3 | ATP-binding cassette sub-family D member 3 | ND |
| 5. | ACAA2 | 3-ketoacyl-CoA thiolase, mitochondrial | MM |
| 6. | ACAD9 | Complex I assembly factor ACAD9, mitochondrial | MIM |
| 7. | ACADVL | Very long-chain specific acyl-CoA dehydrogenase, mitochondrial | MIM |
| 8. | ACAT2 | Acetyl-CoA acetyltransferase, cytosolic | ND |
| 9. | ACBD3 | Golgi resident protein GCP60 | ND |
| 10. | ACOT7 | Cytosolic acyl coenzyme A thioester hydrolase (Isoform 1) | ND |
| 11. | ACSL1 | Long-chain-fatty-acid--CoA ligase 1 | MOM |
| 12. | ACSL3 | Long-chain-fatty-acid--CoA ligase 3 | MOM |
| 13. | ACSL4 | Long-chain-fatty-acid--CoA ligase 4 | MOM |
| 14. | ADH5 | Alcohol dehydrogenase class-3 | ND |
| 15. | AFG3L2 | AFG3-like protein 2 | MIM |

| | | | |
|---|---|---|---|
| 16. | AGPS | Alkyldihydroxyacetonephosphate synthase, peroxisomal | ND |
| 17. | ALDH18A1 | Delta-1-pyrroline-5-carboxylate synthase | MIM |
| 18. | AP3B1 | AP-3 complex subunit beta-1 | ND |
| 19. | ATP5F1A | ATP synthase subunit alpha, mitochondrial | MIM |
| 20. | ATP5F1C | ATP synthase subunit gamma, mitochondrial | MIM |
| 21. | ATP7B | Copper-transporting ATPase 2 | ND |
| 22. | ATPAF1 | ATP synthase mitochondrial F1 complex assembly factor 1 | ND |
| 23. | CAPN1 | Calpain-1 catalytic subunit | ND |
| 24. | CCAR2 | Cell cycle and apoptosis regulator protein 2 | MM |
| 25. | CDK1 | **Cyclin-dependent kinase 1** | ND |
| 26. | CLPX | ATP-dependent Clp protease ATP-binding subunit clpX-like, mitochondrial | ND |
| 27. | COASY | Bifunctional coenzyme A synthase | MM, MOM |
| 28. | DARS2 | Aspartate--tRNA ligase, mitochondrial | MM |
| 29. | DDX1 | ATP-dependent RNA helicase DDX1 | ND |
| 30. | DHX30 | ATP-dependent RNA helicase DHX30 | Mitochondrial nucleoid |
| 31. | DHX36 | ATP-dependent DNA/RNA helicase DHX36 | ND |
| 32. | DLAT | Dihydrolipoyllysine-residue acetyltransferase component of pyruvate dehydrogenase complex, mitochondrial | MM |
| 33. | DNAJC11 | DnaJ homolog subfamily C member 11 | MOM |
| 34. | ECI2 | Enoyl-CoA delta isomerase 2 | ND |
| 35. | ETFA | Electron transfer flavoprotein subunit alpha, mitochondrial | MM |
| 36. | EXD2 | Exonuclease 3'-5' domain-containing protein 2 | MOM, MM |
| 37. | FASTKD2 | FAST kinase domain-containing protein 2, mitochondrial | MM, Mitochondrial nucleoid |
| 38. | FASTKD5 | FAST kinase domain-containing protein 5, mitochondrial | MM, Mitochondrial nucleoid |
| 39. | FLVCR1 | Heme transporter FLVCR1 (mitochondria isoform) | Membrane |
| 40. | GFM1 | Elongation factor G, mitochondrial | MM |
| 41. | GFM2 | Ribosome-releasing factor 2, mitochondrial | MM |
| 42. | GJA1 | Gap junction alpha-1 protein | ND |
| 43. | GLS | Glutaminase kidney isoform, mitochondrial | MM |
| 44. | GPN1 | GPN-loop GTPase 1 | ND |

| | | | |
|---|---|---|---|
| 45. | GUF1 | Translation factor GUF1, mitochondrial | MIM |
| 46. | HADHA | Trifunctional enzyme subunit alpha, mitochondrial | MIM |
| 47. | HAX1 | HCLS1-associated protein X-1 | MM |
| 48. | HEATR1 | HEAT repeat-containing protein 1 | ND |
| 49. | HIP1R | Huntingtin-interacting protein 1-related protein | ND |
| 50. | HSP90AB1 | Heat shock protein HSP 90-beta | ND |
| 51. | IDH3B | Isocitrate dehydrogenase [50] subunit beta, mitochondrial | MM |
| 52. | KARS1 | Lysine--tRNA ligase (Mitochondrial isoform) | MM |
| 53. | KIFBP | KIF-binding protein | ND |
| 54. | KLC2 | Kinesin light chain 2 | ND |
| 55. | LONP1 | Lon protease homolog, mitochondrial | MM |
| 56. | LRPPRC | Leucine-rich PPR motif-containing protein, mitochondrial | Mitochondrial nucleoid |
| 57. | MAPK1 | Mitogen-activated protein kinase 1 | ND |
| 58. | MCCC1 | Methylcrotonoyl-CoA carboxylase subunit alpha, mitochondrial | MM |
| 59. | MFN1 | Mitofusin 1 | MOM |
| 60. | MIPEP | Mitochondrial intermediate peptidase | MM |
| 61. | MRPL38 | Large ribosomal subunit protein mL38 | MIM |
| 62. | MRPL45 | Large ribosomal subunit protein mL45 | MIM |
| 63. | MRPS22 | Small ribosomal subunit protein mS22 | MIM, mitochondrial ribosome |
| 64. | MRPS28 | Small ribosomal subunit protein bS1m | MIM |
| 65. | MSTO1 | Protein misato homolog 1 | MOM |
| 66. | MTCH2 | Mitochondrial carrier homolog 2 | MOM |
| 67. | MTFR1 | **Mitochondrial fission regulator 1** | ND |
| 68. | MTHFD1 | **C-1-tetrahydrofolate synthase, cytoplasmic** | ND |
| 69. | MTOR | Serine/threonine-protein kinase mTOR | MOM |
| 70. | MTPAP | Poly(A) RNA polymerase, mitochondrial | ND |
| 71. | MTX3 | Metaxin-3 | MOM |
| 72. | MUL1 | Mitochondrial ubiquitin ligase activator of NFKB | MOM |
| 73. | NCBP1 | Nuclear cap-binding protein subunit 1 | ND |

| | | | |
|---|---|---|---|
| 74. | NDUFA10 | NADH dehydrogenase [ubiquinone] 1 alpha subcomplex subunit 10, mitochondrial | MIM, MM |
| 75. | NDUFS1 | NADH-ubiquinone oxidoreductase 75 kDa subunit, mitochondrial | MIM, IMS, MM |
| 76. | NOL6 | Nucleolar protein 6 | ND |
| 77. | NRDC | Nardilysin | ND |
| 78. | PC | Pyruvate carboxylase, mitochondrial | MM |
| 79. | PDE2 | cGMP-dependent 3',5'-cyclic phosphodiesterase | MIM, MOM, MM |
| 80. | PEX5 | Peroxisomal targeting signal 1 receptor | ND |
| 81. | PGAM5 | Serine/threonine-protein phosphatase PGAM5, mitochondrial | MOM, MIM |
| 82. | PHB2 | Prohibitin-2 | MIM, MOM |
| 83. | PRKACA | cAMP-dependent protein kinase catalytic subunit alpha | ND |
| 84. | PTCD3 | Small ribosomal subunit protein mS39 | MIM |
| 85. | PYCR1 | Pyrroline-5-carboxylate reductase 1, mitochondrial | MM |
| 86. | PYCR2 | Pyrroline-5-carboxylate reductase 2 | MM |
| 87. | QARS1 | Glutamine--tRNA ligase | MM |
| 88. | QTRT1 | Queuine tRNA-ribosyltransferase catalytic subunit 1 | MOM |
| 89. | RACK1 | Small ribosomal subunit protein RACK1 | ND |
| 90. | RAF1 | RAF proto-oncogene serine/threonine-protein kinase | MOM |
| 91. | RAP1GDS1 | Rap1 GTPase-GDP dissociation stimulator 1 | ND |
| 92. | RARS2 | Probable arginine--tRNA ligase, mitochondrial | MM |
| 93. | RHOT1 | Mitochondrial Rho GTPase 1 | MOM |
| 94. | RHOT2 | **Mitochondrial Rho GTPase 2** | MOM |
| 95. | RPS3 | **Small ribosomal subunit protein uS3** | MIM, MM |
| 96. | SARM1 | NAD(+) hydrolase SARM1 | MOM |
| 97. | SDHA | Succinate dehydrogenase [ubiquinone] flavoprotein subunit, mitochondrial | MIM |
| 98. | SFXN1 | Sideroflexin-1 | MIM |
| 99. | SFXN2 | Sideroflexin-2 | MOM/MIM |
| 100. | SLC25A1 | Tricarboxylate transport protein, mitochondrial | MIM |
| 101. | SLC25A3 | Solute carrier family 25 member 3 | MIM |
| 102. | SLC25A6 | ADP/ATP translocase 3 | MIM |

| | | | |
|---|---|---|---|
| 103. | SLC27A3 | **Long-chain fatty acid transport protein 3** | Membrane |
| 104. | SQSTM1 | Sequestosome-1 | ND |
| 105. | STOML2 | Stomatin-like protein 2, mitochondrial | MIM |
| 106. | SYNE2 | Nesprin-2 | ND |
| 107. | TARS2 | Threonine--tRNA ligase, mitochondrial | MM |
| 108. | TDRKH | Tudor and KH domain-containing protein | ND |
| 109. | TIMM50 | Mitochondrial import inner membrane translocase subunit TIM50 | MIM |
| 110. | TOMM40 | Mitochondrial import outer membrane translocase subunit TOM40 | MOM |
| 111. | TOMM70 | Mitochondrial import outer membrane translocase subunit TOM70 | MOM |
| 112. | TRAP1 | Heat shock protein 75 kDa, mitochondrial | MM, MIM, IMS |
| 113. | TRUB1 | Pseudouridylate synthase TRUB1 | ND |
| 114. | TSFM | Elongation factor Ts, mitochondrial | MM |
| 115. | USP15 | Ubiquitin carboxyl-terminal hydrolase 15 | ND |
| 116. | USP48 | Ubiquitin carboxyl-terminal hydrolase 48 | ND |
| 117. | VARS2 | Valine--tRNA ligase, mitochondrial | ND |
| 118. | VDAC1 | Voltage-dependent anion-selective channel protein 1 | MOM |
| 119. | VDAC2 | Voltage-dependent anion-selective channel protein 2 | MOM |
| 120. | VDAC3 | Voltage-dependent anion-selective channel protein 3 | MOM |
| 121. | YME1L1 | ATP-dependent zinc metalloprotease YME1L1 | MIM |

MM, Mitochondrial matrix; MIM, Mitochondrial inner membrane, IMS, Mitochondrial intermembrane space; MOM, Mitochondrial outer membrane; ND, Sub-mitochondrial localization not determined

**Table S2. Proteins identified from GLUT10-containing vesicles and analyzed by GO enrichment analysis by g:Profiler: CC terms.** A total of 210 identified proteins were analyzed in CC terms.

| Term Name<br>Term ID | – log10<br>Adjusted *P* value | Intersections |
|---|---|---|
| vesicle<br>GO:CC:0031982 | 37.19 | ACTR10,ACTR1B,AGTRAP,AHCY,ALDH7A1,ALDOA,ANP32E,ANXA2,APPL1,ATG9A,ATP5F1B,ATP6V0A2,ATP6V0D1,AZGP1,BCAP31,BPIFB1,BROX,CACYBP,CANX,CAPZA1,CBR1,CCT6A,CD81,CDK1,CFL2,CLIC1,CMTM6,COPA,COPB1,COPB2,COPE,COTL1,CPD,DNAJC7,EIF3H,ENPP4,FAM3C,FSCN1,GDI2,GGCT,GLO1,GOLPH3,GPI,GPR107,GSTK1,HMGB1,HPRT1,HRNR,HSP90AA1,HSPA8,ITM2B,KIAA0319L,M6PR,MPI,MTHFD1,N |

| | | |
|---|---|---|
| | | APA,NEBL,OCLN,PCMT1,PDCD5,PDCD6IP,PDIA3,PIP,PPIB,PRDX1,PTPA,RAB1A,RAB1B,RAB2A,RAB2B,RAB31,RAB35,RAB5B,RAB5C,RAB6A,RAB7A,RAB8A,RABAC1,RARS1,RCC2,RPL12,RPN1,S100A7,SCAMP1,SCAMP2,SCAMP4,SLC12A9,SLC1A5,SLC35F6,SLC38A1,SLC9A8,SNAP29,STMN1,STX10,STX12,STX16,STX6,STX7,STX8,SYAP1,TAGLN2,TCP1,TEX264,TFRC,TGOLN2,TKT,TMEM165,TMEM168,TMEM9,TNPO1,TPI1,TXNDC17,USO1,VAMP2,VAMP3,VAMP4,VAMP8,VCP,VPS35,VTI1A,WDR11,YIPF3,YIPF6,YWHAB |
| endomembrane system<br>GO:CC: 0012505 | 19.12 | ACTR10,ACTR1B,AGTRAP,AIMP1,ALDOA,ANXA2,APPL1,ARFGAP1,ATG9A,ATP2C1,ATP6V0A2,ATP6V0D1,BCAP31,BPNT2,BROX,CACYBP,CANX,CDK1,CLCC1,CLIC1,CMTM6,COPA,COPB1,COPB2,COPE,COTL1,ENPP4,ERLIN1,ERLIN2,FAM3C,GDI2,GOLM1,GOLPH3,GPI,GPR107,GPR108,HMGB1,HRNR,HSP90AA1,HSPA8,ITM2B,KIAA0319L,M6PR,NOSIP,NSF,PARP1,PDCD6IP,PDIA3,PDXDC1,PPIB,RAB1A,RAB1B,RAB2A,RAB2B,RAB31,RAB35,RAB5B,RAB5C,RAB6A,RAB7A,RAB8A,RABAC1,RCC2,RPN1,RTCB,S100A7,SCAMP1,SCAMP2,SCAMP4,SLC35A2,SLC35B2,SLC35C2,SLC9A8,SNAP29,STIP1,STX10,STX12,STX16,STX6,STX7,STX8,SYAP1,TCP1,TEX264,TFRC,TGOLN2,TKT,TMEM115,TMEM165,TMEM168,TMEM87A,TMEM9,TMX1,TPPP,UBIAD1,UNC45A,USO1,VAMP2,VAMP3,VAMP4,VAMP7,VAMP8,VCP,VPS35,VTI1A,WDR11,YIPF3,YIPF4,YIPF6 |
| SNARE complex<br>GO:CC:0031201 | 12.50 | NAPA,SNAP29,STX10,STX12,STX16,STX6,STX7,STX8,VAMP2,VAMP3,VAMP4,VAMP8,VTI1A |
| transport vesicle<br>GO:CC:0030133 | 10.69 | ATP6V0D1,COPA,COPB1,COPB2,COPE,M6PR,RAB1A,RAB1B,RAB5B,RAB6A,RAB7A,RAB8A,RABAC1,SCAMP1,STX12,STX16,STX6,STX7,TGOLN2,TMEM168,USO1,VAMP2,VAMP3,VAMP4,VTI1A,YIPF3 |
| Endosome<br>GO:CC:0005768 | 10.17 | ANXA2,APPL1,ATG9A,ATP6V0A2,ATP6V0D1,CMTM6,GOLPH3,GPR107,HMGB1,HSPA8,ITM2B,M6PR,PDCD6IP,PDIA3,RAB1A,RAB31,RAB35,RAB5B,RAB5C,RAB7A,RAB8A,RCC2,SCAMP1,SCAMP2,SCAMP4,SLC9A8,STX12,STX6,STX7,STX8,TFRC,TGOLN2,TMEM165,TMEM9,VAMP3,VAMP4,VAMP8,VPS35,VTI1A |
| endocytic vesicle<br>GO:CC:0030139 | 9.18 | APPL1,ATP6V0A2,ATP6V0D1,HSP90AA1,M6PR,OCLN,PDIA3,RAB31,RAB35,RAB5B,RAB7A,RAB8A,STX12,STX6,STX7,STX8,TFRC,TGOLN2,VAMP2,VAMP3,VAMP4,VAMP8 |
| clathrin-coated vesicle<br>GO:CC:0030136 | 7.10 | ATP6V0D1,BCAP31,GPR107,HSPA8,M6PR,RAB35,RAB8A,SCAMP1,STX6,TFRC,TGOLN2,VAMP2,VAMP3,VAMP4,VAMP8,VTI1A |
| recycling endosome<br>GO:CC:0055037 | 6.49 | ATG9A,CMTM6,PDIA3,RAB35,RAB8A,SCAMP1,SCAMP2,SCAMP4,STX12,STX6,STX7,STX8,TFRC,VAMP3,VAMP8 |
| early endosome<br>GO:CC:0005769 | 4.98 | ANXA2,APPL1,ATP6V0D1,CMTM6,GPR107,RAB1A,RAB31,RAB5B,RAB5C,RCC2,STX12,STX6,STX7,STX8,TFRC,TMEM165,VAMP3,VAMP8,VPS35 |

**Table S3. YXXΦ-type signals and subcellular compartment targeting.**

| Targeting | Sequence | Protein |
|---|---|---|
| Receptor internalization | [a]4–19-YXXΦ-10–40-Tm | Transferrin receptor, asialoglycoprotein receptor H1, GLUT4 |
| Lysosomal-endosomal membrane | Tm6–9-YXXΦ | CD164, LAMP3 |
| Lysosomal-endosomal membrane (antigen presenting) | Tm6–9-YXXΦ-1 | CD1, CD1b, CD1c, CD1d |
| Intracellular receptor sorting | Tm25–43-YXXΦ-16-132 | CI-MPR, CD-MPR |
| TGN-endosome | Tm26-YXXΦ-4 | TGN38 |
| Endosome | Tm36-YXXΦ-6 | GLUT10 |

[a] Numbers denote amino acid residues before and after the YXXΦ motif. Tm, transmembrane domain; YXXΦ, phenylalanine/tyrosine-based motif; GLUT4, glucose transporter 4; LAMP3, lysosomal-associated membrane protein 3; CI-MPR, cation-independent mannose-6-phosphate receptor; CD-MPR, cation-dependent mannose-6-phosphate receptor; TGN38, *trans*-Golgi network 38. The organization of this table is based on a review by Bonifacino and Traub [26].
